# CDNF rescues human iPSCs-derived dopamine neurons through direct binding to unfolded protein response sensors PERK and IRE1α

**DOI:** 10.1101/2024.12.17.628889

**Authors:** Vera Kovaleva, Olesya Shpironok, Li-Ying Yu, Larisa Ivanova, Satoshi Fudo, Lotta J. Happonen, Urve Toots, Mart Ustav, Tommi Kajander, Mati Karelson, Mart Saarma

## Abstract

Cerebral dopamine neurotrophic factor (CDNF) is an unconventional trophic factor that protects dopamine neurons in cellular and animal models of Parkinson’s disease (PD). CDNF was safe and well tolerated in phase 1 clinical trials for PD treatment, and currently, its peptide analogue is under investigation in phase 1 clinical trials for PD. Despite prominent neuroprotective and neurorestorative activity, the receptors and exact mechanism of CDNF functioning have been obscure. Intracellularly acting CDNF exerts cytoprotection by attenuating endoplasmic reticulum (ER) stress and unfolded protein response (UPR). We demonstrated that this activity occurs through the direct binding of CDNF to ER transmembrane UPR sensors PERK and IRE1α for purified proteins and in cells. We identified CDNF mutants deficient for binding to UPR sensors. CDNF binding to PERK and IRE1α appeared to be crucial for the survival of mouse dopamine neurons in culture. Importantly for clinical translation, CDNF rescues human induced pluripotent stem cell-derived dopamine neurons and promotes their regeneration. CDNF binding to UPR sensors alleviated terminal UPR and promoted neurite outgrowth of human dopamine neurons through direct binding to PERK and IRE1α. CDNF binding to BiP was dispensable for the neuroprotective and neurorestorative activity of CDNF. Therefore, CDNF, or small molecules mimicking its binding to UPR sensors and acting selectively on dopamine neurons with activated UPR, are promising drug candidates for PD treatment.

## INTRODUCTION

Parkinson’s disease (PD) affects nearly 10 million people worldwide, with numbers rising rapidly due to an ageing population. PD is the second most prevalent neurodegenerative disorder after Alzheimer’s disease. It is characterized by the progressive loss of dopamine (DA) neurons in the nigrostriatal pathway, leading to impaired motor function and various non-motor symptoms caused by the degeneration and death of DA neurons in other areas, including the olfactory bulb, cerebral cortex, and enteric nervous system^1–3^. Despite extensive ongoing research, no disease-modifying treatments have been developed to effectively halt the degeneration of DA neurons or regenerate damaged neurons in PD patients^4^. PD drugs alleviating non-motor symptoms have not been described.

Neurodegenerative diseases like PD are associated with ageing and involve disturbances in protein homeostasis, or proteostasis, resulting in the accumulation of misfolded proteins^5^. Impaired proteostasis leads to the endoplasmic reticulum (ER) stress and activates an adaptive signalling machinery called unfolded protein response (UPR). UPR is activated and regulated through three ER transmembrane proteins, UPR sensors, inositol requiring enzyme-1α (IRE1α), activating transcription factor 6 (ATF6), and protein kinase RNA-like endoplasmic reticulum kinase (PERK). Many studies indicate that ER stress and UPR determine the onset and pathology of neurodegenerative diseases caused by protein misfolding^5,6^. The pathogenic role of UPR in PD is unclear.

The histological hallmark of PD is Lewy bodies with α-synuclein as the major component. The misfolding and aggregation of α-synuclein was demonstrated to be accompanied by ER stress^10,11^, and chronic activation of UPR signalling, exacerbating the pathology of the disease^12–15^. Modulating the UPR pathways is neuroprotective in animal models of PD^5,16^, highlighting a new therapeutic avenue for neurodegenerative diseases through restoring protein balance.

The cerebral dopamine neurotrophic factor (CDNF) is an unconventional trophic factor with a unique structure and mode of action^17^. CDNF was shown to be a promising drug candidate for PD^18^. CDNF effectively protected and regenerated DA neurons in various animal models, including the mouse α-synuclein model^19–21^. Recent phase 1 clinical trials in PD provided encouraging results, demonstrating that CDNF is safe and well-tolerated in humans^22^. However, the receptors and precise mechanisms underlying CDNF pro-survival and neuroregenerative action remain to be elucidated.

CDNF is also crucial for neuronal development and survival, as mice deficient in CDNF exhibit age-related loss of enteric neurons, mostly DA neurons, and malfunction of the nigrostriatal dopaminergic system, closely mirroring the early stages of PD^23^.

CDNF is mainly located in the ER, where it modulates the UPR and protects neurons. By regulating the UPR, CDNF mitigated the apoptotic pathways activated in neurodegenerative diseases like PD^17,24,25^. Our recent studies reinforced the notion that the neuroprotective effects of CDNF were mediated by UPR sensor proteins, as inhibiting IRE1α and PERK abolished the neuroprotective action of CDNF from ER stress-induced cell death^26^.

Remarkably, CDNF was shown to interact with α-synuclein. This interaction prevented α-synuclein from entering cells, reduced its phosphorylation, and improved motor function in rodent models of PD, pointing toward a novel approach to mitigating PD pathology^21^. Additionally, CDNF down-regulated pro-inflammatory cytokines and attenuated neuroinflammation, a critical component of PD pathology in mouse models^20,27,28^. The transient transfection of a plasmid encoding human CDNF reduced the 6-hydroxydopamine (6-OHDA)-induced neuroinflammation in the rat substantia nigra^20^. It enhanced the clearance of cellular debris and modulated the immune response, promoting anti-inflammatory and suppressing pro-inflammatory mediators^28^.

Unlike traditional neurotrophic factors that solely function extracellularly, CDNF acts as an extracellular growth factor and an ER-resident protein^17,29^. Quite uniquely, CDNF provides targeted relief to stressed or deteriorating neurons without affecting healthy ones, indicating its specificity in responding to pathological stimuli and clinical safety^26,30^.

We found that CDNF interacts with major ER chaperone BiP (GRP78). However, the binding of CDNF to BiP was not essential for the ability of CDNF to support cell survival^26^. Earlier CDNF was demonstrated to act by regulating UPR signaling^25,26,31,32^, both *in vitro* and *in vivo*^26,33^. Furthermore, research on the paralog of CDNF, mesencephalic astrocyte-derived neurotrophic factor (MANF), shed light on the potential mechanism of action of CDNF. We revealed that the direct interaction of MANF with IRE1α, and not with BiP, is essential for the neuroprotective action of MANF *in vitro* and for protecting DA neurons in an animal model of PD^34^. In the current study, we demonstrated that CDNF can directly bind to all three UPR sensors and regulate their activities. These interactions represent a novel mechanism through which CDNF exerts neuroprotective action in human induced pluripotent stem cell (iPSCs) - derived DA neurons. We also report that CDNF mutants that CDNF mutants that lose binding to UPR sensors are no longer neuroprotective. Thus, we uncover that targeting UPR sensors could trigger survival-promoting neuroregenerative mechanisms in human neurons, offering a potential strategy for treating PD and other ER stress-related diseases.

## RESULTS

### CDNF directly interacts with UPR sensors and BiP abolishes these interactions

Using microscale thermophoresis (MST) and purified proteins we found that CDNF binds directly to all three UPR sensor luminal domains (LDs), with the highest affinity to PERK (K_d_ = 95.3 ± 25.7 nM), then with ATF6 (K_d_ = 146.8 ± 24.7 nM) and with the lowest affinity to IRE1α (K_d_ = 248.1 ± 28.2 nM) (Figures 1A-1C). We demonstrated previously that major ER chaperone BiP interacts with mammalian cell line-produced LDs of UPR sensors with high affinity^34^. Here we found that the binding of CDNF to LDs of UPR sensors was abolished in the presence of 50 nM BiP, indicating that CDNF and BiP share the same or overlapping binding sites in LDs of PERK, IRE1α and ATF6 (Figures 1D-1F). To address whether interaction takes place in a more physiologically relevant setup, we tested the interaction between CDNF and full-length IRE1α using MST for purified CDNF and cell lysates from the cell line expressing IRE1α-green fluorescent protein (GFP) upon doxycycline induction^35^. We found that the affinity of CDNF interacts with full-length IRE1α-GFP was similar to that for the purified LD of IRE1α protein (Figure 1G).

**Figure 1.**
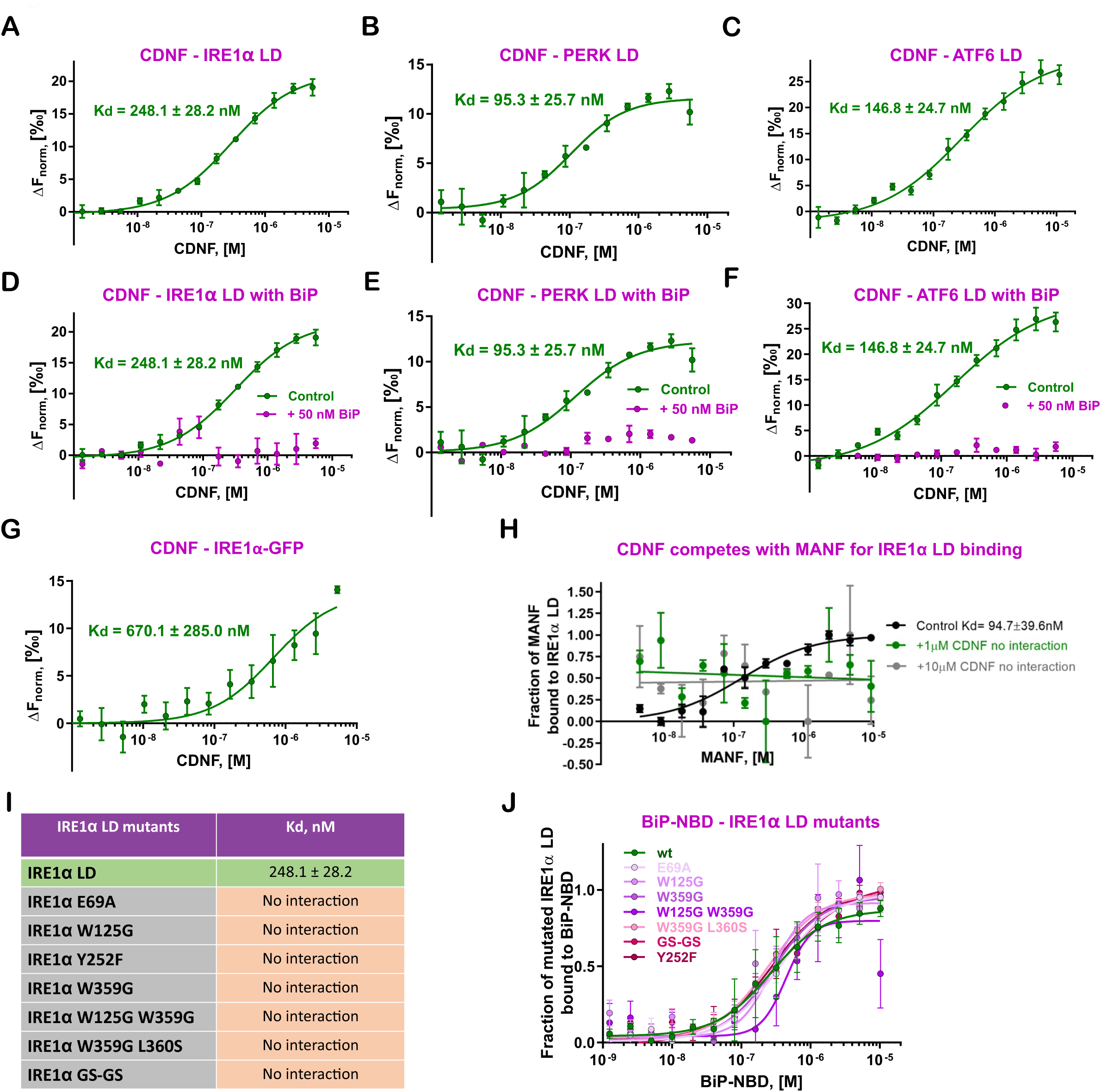
CDNF directly interacts with UPR sensors and competes with BiP/unfolded protein and MANF for binding as shown using microscale thermophoresis (MST). (A), (B), (C) Purified recombinant CDNF protein (0–5.5 µM) interacts with IRE1α, PERK, and ATF6 luminal domains (LDs) labeled through His-tag (20 nM). (D), (E), (F) Interaction of CDNF protein (0–5.5 µM) with IRE1α, PERK, ATF6 LDs (20 nM) labeled through His-tag is abolished in the presence of BiP (50 nM). (G) CDNF (0–5.3 µM) interacts with full-length IRE1α-GFP in crude cell lysates. (H) MANF (0–9.3 µM) interaction with IRE1α LD (20 nM) is abolished in the presence of 1 µM and 10 µM CDNF. (I) Purified recombinant CDNF protein (0–5.5 µM) does not interact with IRE1α mutants (all at 20 nM) labeled through His-tag. In IRE1α GS-GS (oligomerization-deficient mutant)^70^, WLLI_359-362_ were mutated to GSGS_359-362_. (J) Purified recombinant BiP nucleotide-binding domain (NBD) (0–5.1 µM) interacts with all IRE1α mutants (all at 20 nM) with similar affinities to that for wt IRE1α. In (A) – (J) MST binding curves, showing mean fraction bond values from n = 3–5 experiments per binding pair ± SEM, K_d_ values ± error estimations are indicated.

### CDNF competes with MANF for IRE1α binding

Since both MANF and CDNF interact with LDs of UPR sensors and their interaction with UPR receptors is abolished in the presence of BiP, and they share structural similarities, we hypothesized that CDNF may affect the binding of MANF to UPR sensors. We tested how the presence of CDNF affects the interaction of MANF with IRE1α LD. In the presence of 1 µM or 10 µM CDNF, this interaction was abolished, suggesting a similar binding mode for IRE1α LD binding for these homologous proteins (Figure 1H). To test whether the binding of CDNF to IRE1α LD mutants deficient for MANF binding is compromised, we used the IRE1α LD mutants generated in our previous study^34^. We found that all the IRE1α mutants, including IRE1α W125G able to bind MANF, were deficient for CDNF binding (Figure 1I). To ensure that IRE1α LD mutants are active and specifically deficient for CDNF binding, we tested their interaction with a known binding partner, the nucleotide-binding domain of BiP (BiP NBD). The affinities of BiP NBD to IRE1α LD mutants were like that of wild type IRE1α LD (Figure 1J). These data demonstrated that CDNF and MANF binding sites on IRE1α are overlapping but not identical, as IRE1α W125G mutation did not affect the binding of IRE1α to MANF, but abolished that to CDNF^34^.

### CDNF interacts with IRE1α and PERK in cells

To test whether the interactions of CDNF with PERK and IRE1α occur in cells we performed an *in situ* proximity ligation assay (PLA) using Flp-In-T-REx 293 cells. The cell lines were engineered to express either CDNF-hemagglutinin (HA) or GFP-HA driven by a doxycycline-inducible promoter^26^. The PLA results support that biologically active CDNF-HA interacts with endogenous PERK and IRE1α (Figure 2A and 2B). The interactions of GFP with IRE1α or PERK in corresponding cell lines served as negative control. The number of interaction events for CDNF-PERK binding was twofold higher as compared with that for IRE1α, which aligns with our MST data indicating that PERK has a higher affinity for CDNF and therefore may be its primary receptor (Figure 2B).

**Figure 2.**
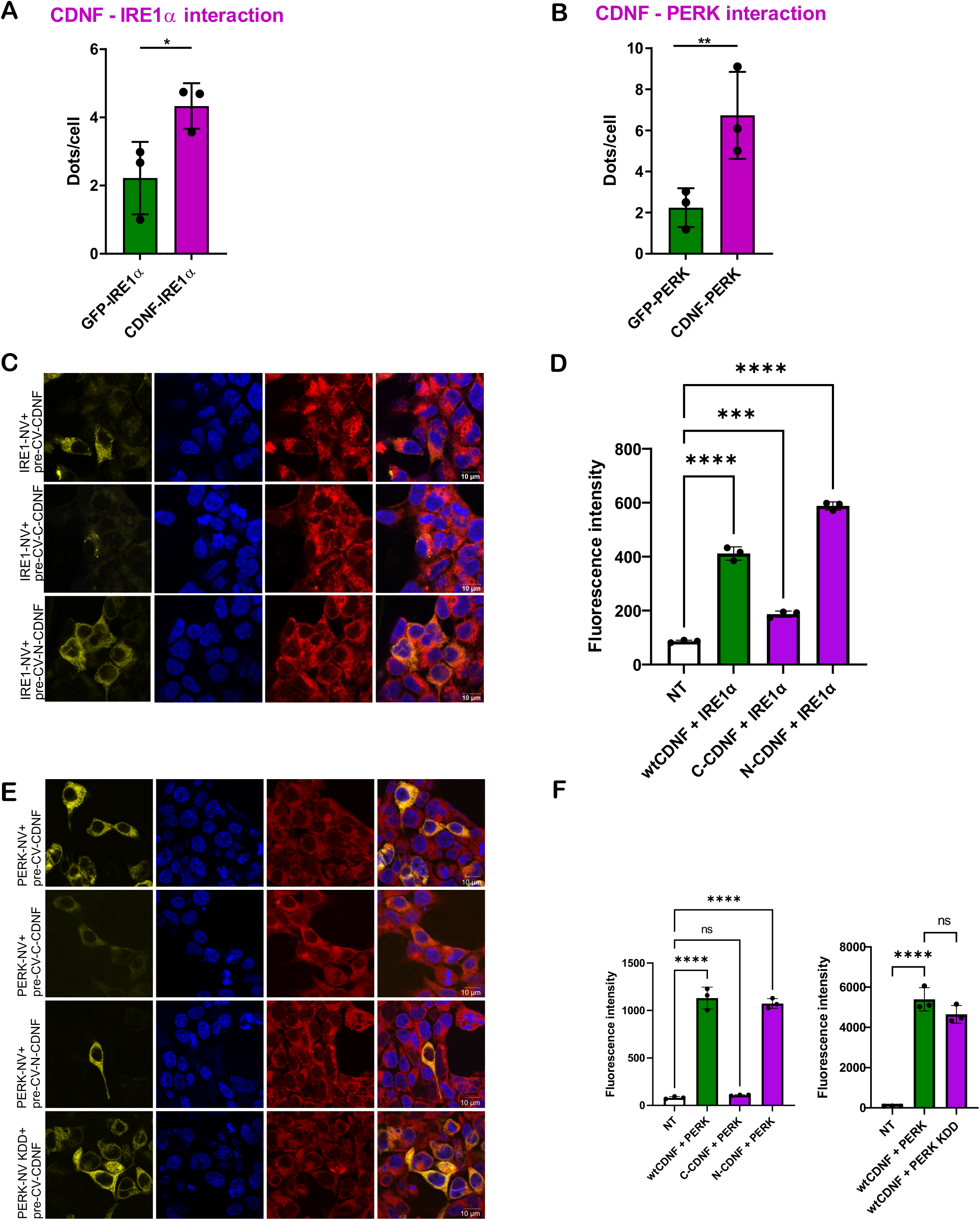
CDNF interacts with IRE1α and PERK in cells. (A) Quantification of CDNF interaction with IRE1α in HEK293T cells, shown as dots/cell, using proximity ligation assay. Mean dots/cell values ± SEM from n = 3 independent experiments are indicated. (B) Quantification of CDNF interaction with PERK in HEK293T cells, shown as dots/cell, using proximity ligation assay. Mean dots/cell values ± SEM from n = 3 independent experiments are indicated. (C) CDNF, C-terminal domain of CDNF (C-CDNF), N-terminal domain of CDNF (N-CDNF) interaction with IRE1α and in HEK293 cells in bimolecular fluorescence complementation (BiFC) assay. A yellow, fluorescent signal indicates interaction between the tested proteins. The endoplasmic reticulum (ER) was visualized using ER-ID-red dye (red) and nuclei by Hoechst 33342 staining (blue). (D) Quantification of fluorescence intensity in IRE1α BiFC assays. Cells expressing wild-type CDNF (wt CDNF), C-terminal CDNF (C-CDNF), or N-terminal CDNF (N-CDNF) were co-transfected with IRE1α and analyzed by flow cytometry. The fluorescence intensities of live cells were measured using an LSRFortessa flow cytometer with a 100 µm nozzle and 488 nm excitation laser. A total of 10,000 live cells were evaluated in the GFP channel (527/32 nm bandpass filter) after gating to exclude debris and non-uniform cells. Shown are representative images from n = 3–5 independent experiments per binding pair. Mean ± SD; n = 3, ordinary one-way ANOVA with Dunnett’s test, ***p<0.001, ****p<0.0001 (E) CDNF, C-terminal domain of CDNF (C-CDNF), N-terminal domain of CDNF (N-CDNF) interaction with PERK and in HEK293 cells in bimolecular fluorescence complementation (BiFC) assay. A yellow, fluorescent signal indicates interaction between the tested proteins. Endoplasmic reticulum (ER) was visualized using ER-ID-red dye (red) and nuclei by Hoechst 33342 staining (blue). (F) Left, Quantification of fluorescence intensity in PERK BiFC assays. Cells co-expressing PERK and wild-type or mutant CDNF (C-CDNF, N-CDNF, or KDD PERK mutant) were analyzed similarly as in panel D. Fluorescence readout was performed by flow cytometry as described, using an OD600=0.5 cell culture sample. 10,000 live cells were evaluated, and fluorescence intensities were measured using the GFP channel. Statistical analysis was performed using one-way ANOVA with Dunnett’s test, with significance levels indicated as follows: ns = not significant, ****p<0.0001. Right, Cells co-expressing PERK with wild-type CDNF (wt CDNF), C-CDNF, N-CDNF, or the PERK KDD mutant were analyzed. The interaction of wt CDNF and N-CDNF with PERK has significantly increased fluorescence intensity, indicating strong interaction. C-CDNF did not result in a significant increase in signal, indicating that its interaction with PERK is weak or diminished. No difference in fluorescence intensity was observed between PERK KDD and wild-type PERK when co-expressed with wt CDNF, suggesting that the PERK KDD mutation does not affect the interaction of CDNF with PERK.

We further analysed CDNF interaction with PERK and IRE1α in HEK293 using a bimolecular fluorescence complementation (BiFC) assay. We generated constructs for the CDNF, PERK, and IRE1α with non-fluorescent fragments of C-Venus (CV) and N-Venus (NV) in the same vector and under the same promotor for all constructs, as described earlier in^34^. The fluorescent signals were quantified using flow cytometry. To validate the BiFC signal generation, we used two pairs of interacting proteins, transcription factors Jun-NV and Fos-CV, and ER-located BiP-NV and PERK-CV, as positive controls. The interactions of these protein pairs were confirmed previously and were specifically chosen to validate the occurrence of BiFC signals within the nucleus and the ER, respectively (Figure S2C and S2C).

We detected the fluorescent signal for CDNF-IRE1α and CDNF-PERK combinations (Figures 2C‒2F). Importantly, the fluorescence intensity of CDNF binding to PERK was 7 times higher than that observed in the CDNF-IRE1α interaction. Although these differences can be partially explained by a higher level of PERK expression, compared to that of IRE1α (Figure S2D) they further support the notion that PERK is the primary receptor for CDNF.

CDNF consists of the lipid-binding N-terminal domain (N-CDNF) and the C-terminal domain (C-CDNF) carrying cytoprotective activity^26,36^. The two domains are connected by a flexible linker region, and we have shown that the domains can function independently^13,20^. C-CDNF contains a disulfide-forming CXXC loop, a motif typical for oxidoreductases^37^, and an ER retention sequence (KDEL), carrying out the retrieval of ER proteins from the Golgi, preventing the protein secretion to the extracellular space. To address the role of CDNF domains in the binding to PERK and IRE1α we tested the interaction between N-terminal and C-terminal domains of CDNF with PERK and IRE1α in the BiFC assay. Both the C-CDNF and N-CDNF constructs contained signal sequences. Given the absence of an ER retention signal in the N-CDNF construct, it is likely more readily secreted than full-length CDNF. Remarkably, the fluorescent signal for the N-CDNF was more evenly distributed in the ER than that for the C-CDNF and full-length CDNF (Figure 2C). In microscopy imaging, we observed different fluorescent signals for C-CDNF and N-CDNF. Therefore, we quantified the intensities of the signal for CDNF domains and PERK and IRE1α binding using flow cytometry. The fluorescent signal for C-CDNF was significantly lower while the signal for N-CDNF was similar to that of full-length CDNF (Figure 2D). There was no statistically significant difference in the fluorescence intensity between full-length CDNF and N-CDNF, indicating that the N-terminal domain of CDNF also contributes to the binding of CDNF with PERK. In contrast, the C-terminal domain (C-CDNF) demonstrated a significantly weaker interaction with PERK (Figures 2E and 2F).

The expression level of C-CDNF is about 2 times lower and that of N-CDNF is slightly lower than CDNF (though not statistically significant) (Figure S2E). Therefore, we can conclude that C-CDNF and full-length CDNF bind to IRE1α with similar strength, and the N-terminus of CDNF may have stronger binding to IRE1α. For PERK, the BiFC data suggest the binding of full-length CDNF and N-CDNF are similar and C-CDNF appears to be weaker.

We further aimed to understand if the kinase activity of PERK plays a role in its interaction with CDNF. To explore this, we investigated the interaction of CDNF and a kinase-dead mutant of PERK (PERK KDD)^38^. Our data indicate that CDNF interacts with PERK and PERK KDD demonstrating that the active kinase domain is not crucial for this interaction. The intensities of the fluorescent signals from the interactions of CDNF with PERK and its kinase-dead mutant (PERK KDD) were comparable (Figures 2E and 2F).

### Identification of CDNF regions involved in binding to PERK

The binding between CDNF and PERK LD was modelled using protein-protein docking, similarly to our previous work^34^. For the identification of the possible PERK binding sites in CDNF, the available three-dimensional (3D) structures of CDNF (PDB ID: 4BIT)^39^ and PERK LD (PDB ID: 4YZS)^40^ were used. The protein-protein docking results predicted the possible CDNF mutants deficient in PERK binding.

The protein-protein docking procedure was carried out without any mutual protein position constraints. The CDNF, as a ligand, was docked flexibly and PERK LD was considered a receptor. Analysis of the interactions between CDNF and PERK LD in 30 computationally generated models of the CDNF – PERK LD complexes suggested that CDNF has two possible PERK LD binding sites. The first possible binding site is in the C-terminal domain of CDNF. It involves the conserved cysteine loop ^132^CRAC^135^. The second possible PERK LD binding site on CDNF is located partly in the linker region between the C- and N-terminal domains of CDNF and partly in the N-terminus of CDNF, it involves residues K85– Y102, and residues R54–D64, respectively (Figure 3B). The linker region CDNF binding site overlaps with MANF binding sites to IRE1α^34^ and is similar to the second possible binding site on CDNF. Further analysis of the docking results was conducted as described in our previous work^34^. Briefly, the importance of individual amino acid residues of CDNF in the interaction with PERK LD was calculated based on the frequency of each amino acid residue’s appearance and the number of specific interactions (*i.e.,* hydrogen bonds) in 30 docking models. This analysis allowed us to identify several amino acid residues on both predicted possible binding sites of CDNF that have the most frequent involvement in specific interactions. For the first possible binding site, the K122, Q123, E130, E131, and R133 amino acid residues were suggested as the most probable candidates for PERK interaction and consequently for point mutations (Table 1). For the second possible binding site, amino acid residues Y57, Y58, K63, K85, K89, S95, Q96, and Y102 were proposed as targets for mutations (Table 1).

**Figure 3.**
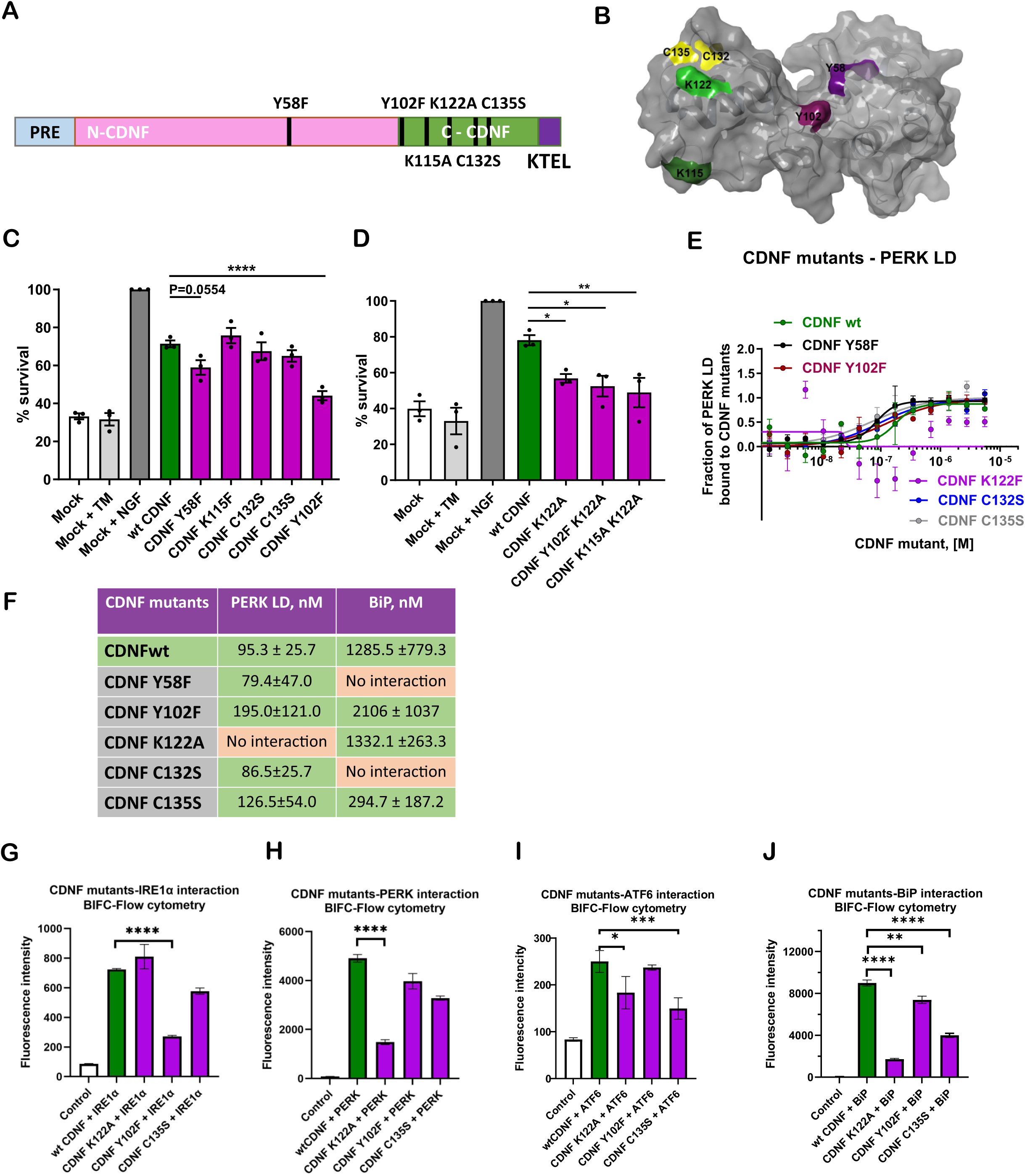
CDNF mutants deficient for binding to UPR sensors were generated and characterized. (A) Scheme of two-domain CDNF structure with sites of mutation. (B) A structure of CDNF with mutated amino acids indicated in yellow, green and purple. (C) Microinjections of CDNF Y58F, CDNF Y102F encoding plasmids to SCG neurons do not rescue them from tunicamycin (TM) -induced cell death compared with mock-injected with empty vector or uninjected. TM was used at 2 µM for 72 hours. Mean ± SEM; n = 3. (D) Microinjections of CDNF K122A, CDNF Y102F K122A and CDNF K115A K122A encoding plasmids to SCG neurons do not rescue them from TM-induced cell death compared with mock-injected with empty vector or uninjected. Mean ± SEM; n = 3. (E) Interactions of CDNF mutants with PERK LD labeled through His-tag. MST binding curves, showing mean fraction bond values from n = 3–5 experiments per binding pair ± SEM. Kd values ± error estimations are indicated. (F) Binding affinities (Kd) of CDNF mutants binding to PERK LD and BiP. (G, H, I, J) BiFC signal of the interactions of CDNF mutants with IRE1α, PERK, ATF6 and BiP in HEK293 cells. Shown are representative images from n = 3–5 independent experiments per binding pair. Mean ± SD; ordinary one-way ANOVA with Dunnett’s test, *: p ≤ 0.05; **: p ≤ 0.01. (Heat-map for mutants Kd)

**Table 1.** The analysis of the frequency of the appearance of each interacting amino acid residues in computationally generated models of the CDNF–PERK LD complexes 1-30.

| CDNF amino acid residue | Number of models in which the amino acid residue is involved in binding to PERK LD |  |  |
| --- | --- | --- | --- |
|  | Total | Total % | Hydrogen bonds |
| Binding site 1 |  |  |  |
| K122 | 10 | 33.3 | 10 |
| Q123 | 10 | 33.3 | 7 |
| L125 | 7 | 23.3 | 0 |
| H126 | 20 | 66.7 | 7 |
| S127 | 16 | 53.3 | 3 |
| T128 | 15 | 50.0 | 1 |
| G129 | 23 | 76.7 | 3 |
| E130 | 18 | 60.0 | 7 |
| E131 | 21 | 70.0 | 8 |
| C132 | 9 | 30.0 | 0 |
| R133 | 17 | 56.7 | 10 |
| A134 | 6 | 20.0 | 0 |
| C135 | 4 | 13.3 | 0 |
| **Binding site 2** |  |  |  |
| R54 | 9 | 30.0 | 0 |
| Y57 | 21 | 70.0 | 10 |
| Y58 | 22 | 73.3 | 5 |
| K63 | 6 | 20.0 | 3 |
| D64 | 14 | 46.7 | 4 |
| K85 | 6 | 20.0 | 3 |
| K89 | 6 | 20.0 | 6 |
| K91 | 5 | 16.7 | 2 |
| K92 | 7 | 23.3 | 1 |
| L93 | 17 | 56.7 | 0 |
| D94 | 14 | 46.7 | 4 |
| S95 | 10 | 33.3 | 4 |
| Q96 | 21 | 70.0 | 10 |
| Y102 | 18 | 60.0 | 6 |

### Validation of CDNF binding sites to PERK

Using molecular docking, we identified potential binding sites implicated in the interactions between CDNF-PERK and predicted specific CDNF mutations to verify these binding sites and analyse their functional implications. We engineered eight plasmid vectors carrying CDNF mutants targeting critical regions believed to be pivotal for its interaction with PERK. These mutants include Y58F in the N-terminal domain of CDNF, and Y102F, K115A, K122A, C132S and C135S in the C-terminal domain (Figures 3A and 3B). Furthermore, we developed two double mutants, Y102F-K122A and K115F-K122A so that a few potential PERK-binding sites in CDNF could be affected simultaneously.

To examine the effect of mutations on neuronal survival-promoting activity of CDNF the plasmids encoding CDNF mutants were microinjected into early postnatal mouse superior cervical ganglion (SCG) sympathetic neurons. Following this, the neurons were exposed to the ER stressor tunicamycin (TM) triggering apoptosis of SCG neurons. Our earlier data demonstrate that CDNF is neuroprotective in this model against TM-induced cell death^26^. We found that CDNF with mutations Y102F and K122A, and the double mutations Y102F K122A and K115A K122A, was not able to protect TM-treated SCG neurons, and the Y58F mutation in the N-terminal region of CDNF attenuated neuronal survival, as compared with wt CDNF (Figures 3C and 3D). In contrast, the K115A, C132S, and C135S CDNF mutants were biologically active, and these mutations did not significantly affect neuronal survival and neuroprotection by CDNF (Figure 3C and 3D). Plasmid microinjection of SCG neurons with CDNF mutants Y58F, Y102F, and K122A resulted in reduced neuroprotective activity, while the C132S and C135S mutants remained biologically active. To verify that the observed phenotype correlates with the ability to interact with PERK, we produced recombinant proteins for these CDNF mutants. Since the CDNF double mutants contain a mutation in position K122A that reduces its biological activity and may impair the neuroprotective function of double mutants, we chose to produce only the K122A mutant and not K122A-containing double mutants, as proteins for further investigations.

To assess the impact of specific amino acid residues on the interaction network of CDNF, we used MST to analyse the binding affinities of five CDNF mutants with PERK LD. The CDNF mutants Y58F, Y102F, C132S, and C135S retained their binding ability for the PERK LD with affinities and K_d_ values comparable to that of wt CDNF, whereas CDNF K122A was deficient for PERK binding (Figures 3E and 3F). Additionally, we tested the CDNF mutant proteins for binding to the ER chaperone BiP, demonstrated to form a complex with CDNF^26,41^. We found that all mutants except for Y58F and C132S interacted with BiP, and cysteine loop mutants C135S had higher affinity than wt CDNF (Figure 3F).

We further investigated the interactions of these mutants using BiFC assay. We created constructs of the CDNF mutants with non-fluorescent fragments of C-Venus and N-Venus and then tested their ability to bind to PERK, IRE1α, ATF6 and BiP. The intensity of the BiFC fluorescence generally reflects the strength of the protein-protein interaction, but differently from MST where two purified proteins interact, in BiFC other cellular proteins may influence the interaction. Intriguingly, for IRE1α, the fluorescent signal observed for the CDNF Y102F was almost abolished, and for CDNF C135S it was weaker, but for CDNF K122A it was similar to that for wt CDNF (Figure 3G). We further tested the interaction of CDNF mutants with PERK and found that the fluorescent signal for the CDNF K122A mutant binding was 5 times lower than that for both wt CDNF and the other mutants (Figure 3H). We also assessed the interaction wt CDNF and CDNF mutants with ATF6 using BiFC and found that the interaction of CDNF C135S was significantly reduced, the fluorescent signal for CDNF K122A was slightly reduced, and for CDNF Y102F, it was not affected (Figure 3I) The fluorescent signal of the interaction between the CDNF K122A mutant and BiP was significantly reduced, the signal for CDNF C135S interaction with BiP was also reduced (Figure 3J).

### Crosslinking mass spectrometry analysis suggests a binding mode for CDNF to PERK LD

To further validate the interactions between CDNF and PERK, we performed crosslinking mass spectrometry on purified recombinant complexes using disuccinimidyl suberate (DSS). Based on the interprotein crosslinks identified, we docked the protein complex in two distinct phases using the HADDOCK platform^42^. An AlphaFold2 model of the PERK LD was generated to fill in missing loops from the crystal structure, providing a complete model for molecular docking studies. For CDNF we used the available full-length NMR ensemble structure^39^ for the modelling of the CDNF-PERK complex. In the first phase, we generated a model incorporating all identified interprotein crosslinks, selecting the best cluster for further analysis. In the second phase, we excluded one outlier crosslink^43^ and any crosslinks to the PERK LD N-terminus. This exclusion was due to the challenge of reliably predicting the position of the flexible N-terminus^40,44^. Despite this, all crosslinks to the PERK N-terminus were within the allowed 30Å cutoff distance for DSS^45^. The final model was selected based on the highest HADDOCK score and a detailed manual inspection of the interface. Our results indicate that CDNF interacts with PERK in a manner consistent with the crosslinking data and protein-protein-docking predicted models. The flexible nature of the PERK N-terminus presents a challenge for precise modelling, but our approach ensured that all CDNF-PERK LD interactions fell within acceptable distance thresholds. The findings suggest that the α-helix 118-128 and the loop region 129-134 of the C-CDNF would interact with the PERK LD surface involving the N-terminal β-sheet of the PERK LD in residue range 114-133 and region 204-234, involving 3 short β-strands and loops, and a short helical region near the PERK LD dimer interface, but on opposite side of the latter β-sheet (Figure 4). Importantly, the model is also consistent with K122 of CDNF as a key residue for the interaction, as it is crosslinked directly to the interface (Figure 4, Figure S4).

**Figure 4.**
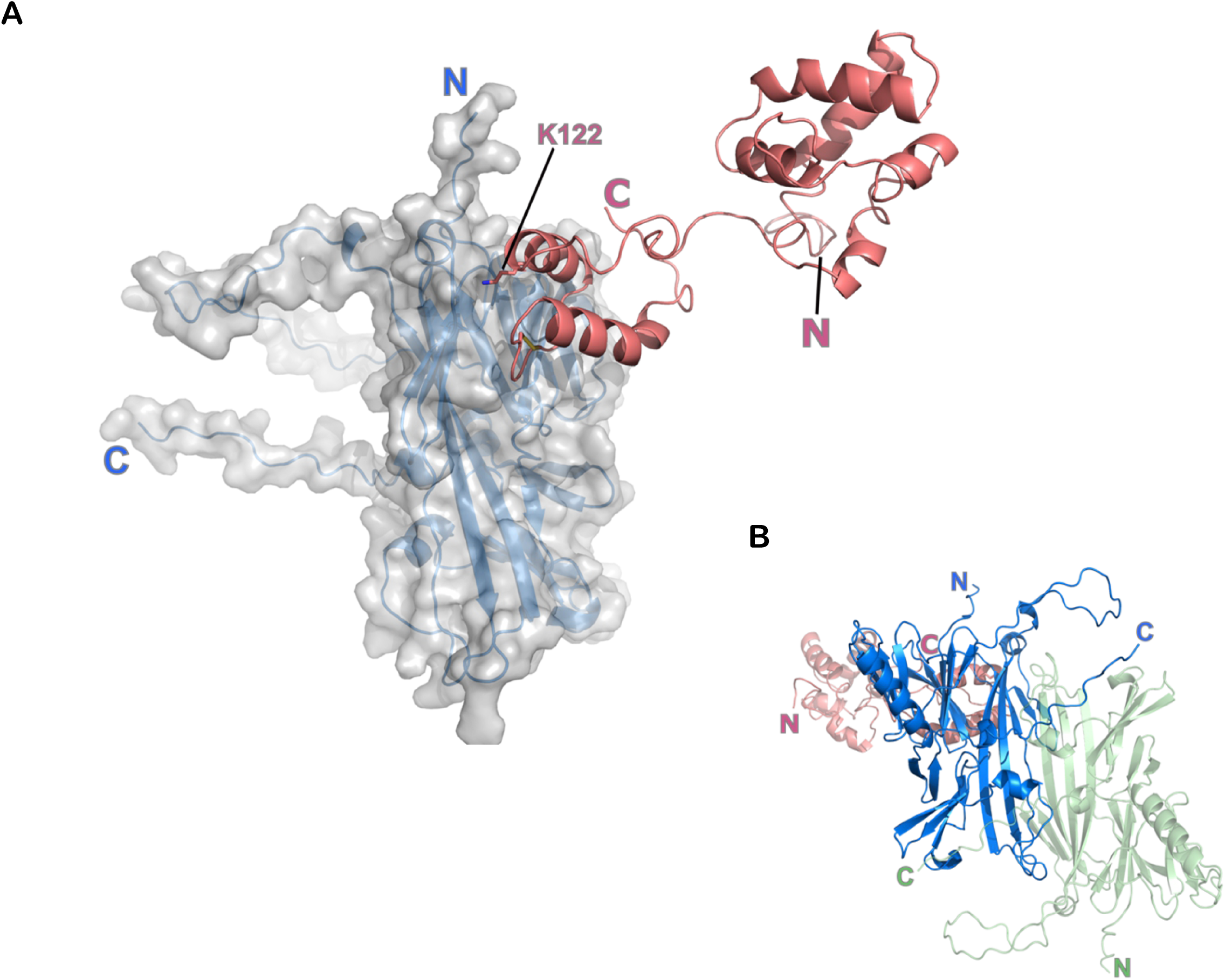
Model for CDNF-PERK complex structure. (A) A model of CDNF-PERK complex structure from molecular docking and modelling based on crosslinking distance constraints. The PERK luminal domain is shown as a transparent surface and blue cartoon, CDNF as a red cartoon, and CDNF Lys122 and the C-CDNF disulfide shown with sticks. “N” and “C” mark the N and C-termini of the proteins. (Insert) The complex in the context of the PERK LD dimer in blue and green for each monomer. View looking down the 2-fold axis of PERK LD dimer, CDNF in red.

### CDNF regulates UPR-related genes upon ER stress

Severe ER stress and hyperactivated UPR can lead to inflammation and apoptosis, and trigger the pathophysiological processes in neurodegenerative diseases^5,9^, CDNF protects and restores the function of dopamine (DA) neurons through the attenuation of UPR signalling^13^. To understand how CDNF rescues the neurons through UPR sensors and regulates UPR-induced apoptosis, we analysed, how exogenously added CDNF and CDNF mutants affect the expression of UPR-related genes. CDNF is expressed in a wide range of neuronal and non-neuronal tissues, though its abundance is generally low. Although a high concentration of CDNF is found in skeletal muscle, heart and testis, MANF’s expression levels in other tissues and cell lines are approximately 100 times higher than those of CDNF^46^ (data also available from the Human Protein Atlas, https://www.proteinatlas.org/). We used the human multiple myeloma RPMI8226 cell line as a model since it has a high expression level of endogenous CDNF. The cells were treated with TM, inhibiting N-glycosylation to induce ER stress, and then recombinant wt CDNF protein or CDNF mutants K122A and Y102F at 100 ng/ml were added to the media and incubated for 72 hours.

As expected, treatment with TM significantly increased the mRNA levels of UPR genes, *sXBP1,* induced through IRE1α, *ATF4* and *CHOP*, induced through PERK and *ATF6* and *BiP*, induced through ATF6, by 2-4-fold (Figure 5A–F). CDNF applied together with TM significantly reduced the expression of *ATF4*, but did not affect *sXBP1, BIP*, or *CHOP* transcript levels (Figure 5A and Figure S5A-S5I). CDNF K122A was not affecting the mRNA level of *ATF4*, whereas CDNF Y102F decreased them similarly to wt CDNF (Figure 5A).

**Figure 5.**
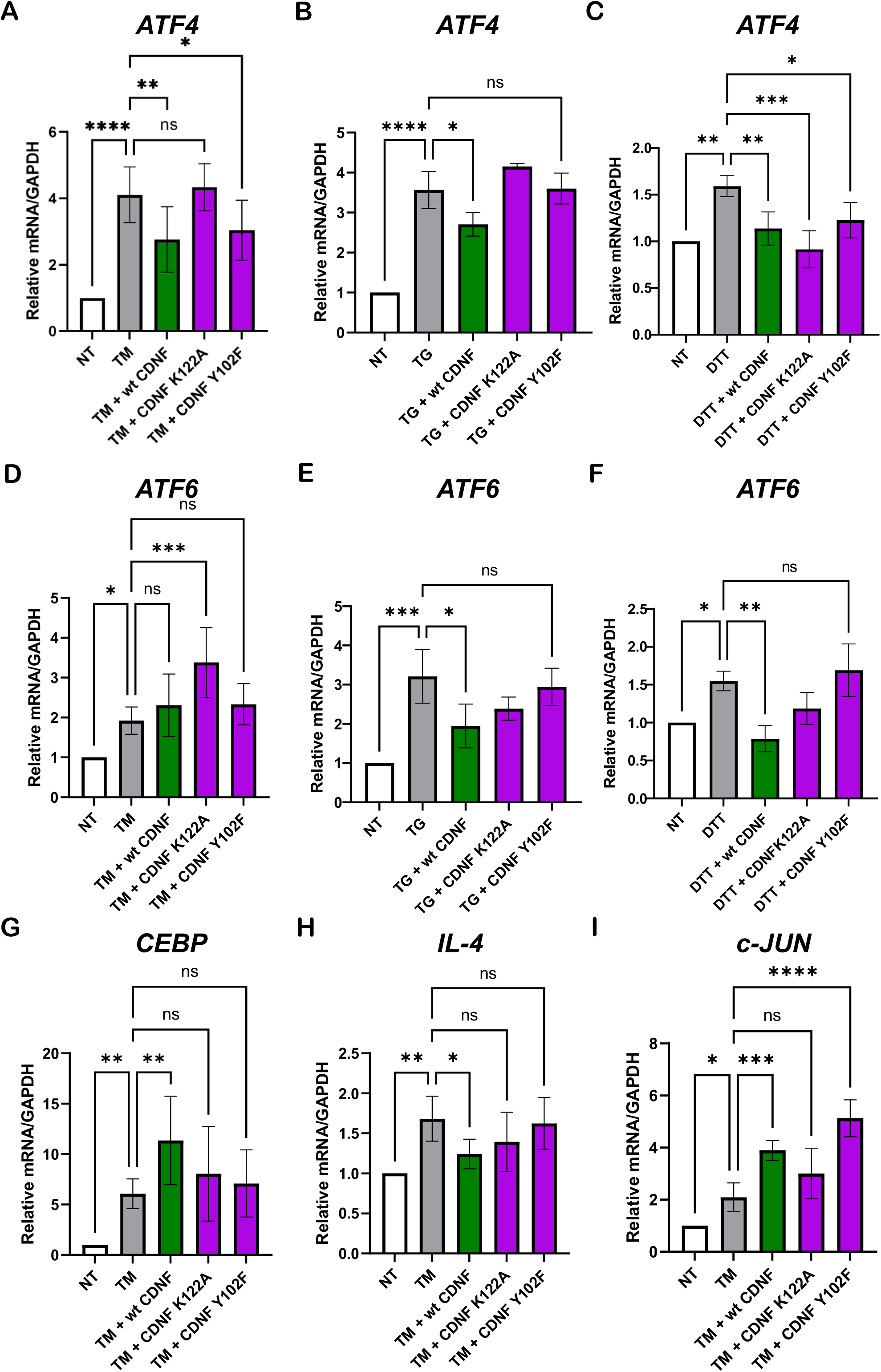
CDNF regulates UPR signalling and inflammation genes *in vitro* through binding to IRE1α, PERK and ATF6. (A‒I) *ATF4, ATF6, CEBPB, IL-4* and *C-JUN,* mRNA levels in RPMI8226 cells. RPMI8226 cells were cultured 24 hours before inducing ER stress by tunicamycin (TM), thapsigargin (TG) or DL-Dithiothreitol (DTT). Cells were treated for 4 hours with TM (2.5 µg/ml), TG (1 µM), or DTT (2 mM), followed by 20 hours of treatment with 400 nM CDNF, CDNF K122A, or CDNF Y102F. The expression levels of UPR marker transcripts were determined by RT-qPCR. Each transcript’s levels have been normalized to the *GAPDH* housekeeping gene and presented as a fold change to the respective control data set from non-treated cells. Shown are means of n = 3–6 experiments ± SD, ordinary one-way ANOVA with Dunnett’s test, *: p ≤ 0.05; **: p ≤ 0.01.

To analyse if the mechanism of ER-stress induction affects the efficacy of CDNF, we tested whether the mRNA levels of UPR-related genes are regulated upon the treatments with dithiothreitol (DTT) and thapsigargin (TG), inducing ER-stress through the disruption of disulfide bond formation in proteins, and block of the ER Ca²⁺-ATPase (SERCA), respectively^47^. Our results demonstrated that CDNF downregulated the expression of *ATF4* in the presence of TG, whereas both CDNF Y102F and CDNF K122A lacked this regulatory effect on the mRNA level of *ATF4* (Figure 5B). However, upon the reductive stress induced by DTT, both CDNF and CDNF Y102F and CDNF K122A downregulated *ATF4* (Figure 5C).

CDNF and CDNF Y102F did not affect the mRNA levels of *ATF6*, while CDNF K122A upregulated it (Figure 5D). In contrast, upon TG- and DTT-induced ER stress CDNF reduced the mRNA level of *ATF6*, whereas both CDNF Y102F and CDNF K122A did not affect it in these conditions (Figures 5E and 5F). Upon reductive stress with DTT, the regulation of *ATF6* was in line with TG treatment. Consistent with our findings, mutations in the CXXC motif of CDNF do not abolish its ability to support the survival of SCG neurons treated with TM. Additionally, as we showed earlier^26^, modifying the CXXC motif in CDNF, specifically through the CDNF C135S mutation, decreases, but does not eliminate CDNF’s ability to promote neuronal survival upon TM treatment. This implies that fully inactivating the CXXC bridge does not affect its neuroprotective role against apoptosis induced by ER stress in SCG neurons. In addition, we tested whether CDNF or CDNF mutants affect the level of *BIP*, *CHOP*, and *sXBP1* transcripts. No significant changes in the mRNA levels of these UPR markers were found (Figure S5). Since CDNF significantly reduces inflammation *in vivo*^20,28^, we further examined whether CDNF could influence the expression of genes associated with the immune response and inflammation. We found that CDNF upregulated the mRNA level of CCAAT/enhancer binding protein (CEBP), mediating the levels of anti-inflammatory and pro-inflammatory genes during the tissue repair (Figure 5G), downregulated the mRNA level of anti-inflammatory interleukin-4 (IL-4) (Figure 5H) and upregulated the mRNA level of *c-JUN*, regulating the activation of macrophages (Figure 5I). CDNF Y102F and CDNF K122A lost the ability to regulate *CEBP* and *IL-4* mRNA levels (Figures 5G and 5H). CDNF K122A did not affect the *c-JUN* mRNA level, whereas the upregulation of the *c-JUN* mRNA level by CDNF Y102F was higher, as compared with that by CDNF (Figure 5I).

### CDNF protects mouse sympathetic neurons through binding to PERK and IRE1α

We further tested whether the deficiency in binding to UPR sensors PERK and IRE1α affects the neuroprotective activity of CDNF in mouse sympathetic SCG neurons when CDNF and its variants were microinjected as purified proteins, upon TM-induced ER stress. We found that IRE1α binding deficient CDNF Y58F and CDNF Y102F were not rescuing these neurons from ER stress-induced cell death triggered by tunicamycin, as efficiently as wt CDNF (Figure 6A). Similarly to IRE1α binding deficient mutants, CDNF K122A deficient for PERK binding did not protect SCG neurons either, indicating that both PERK and IRE1α binding determine the pro-survival and neuroprotective activity of CDNF in mouse SCG neurons (Figure 6A). In contrast, cysteine loop mutants CDNF C132S and CDNF C135S were protecting mouse SCG neurons from ER stress similarly to wt CDNF (Figure 6A).

**Figure 6.**
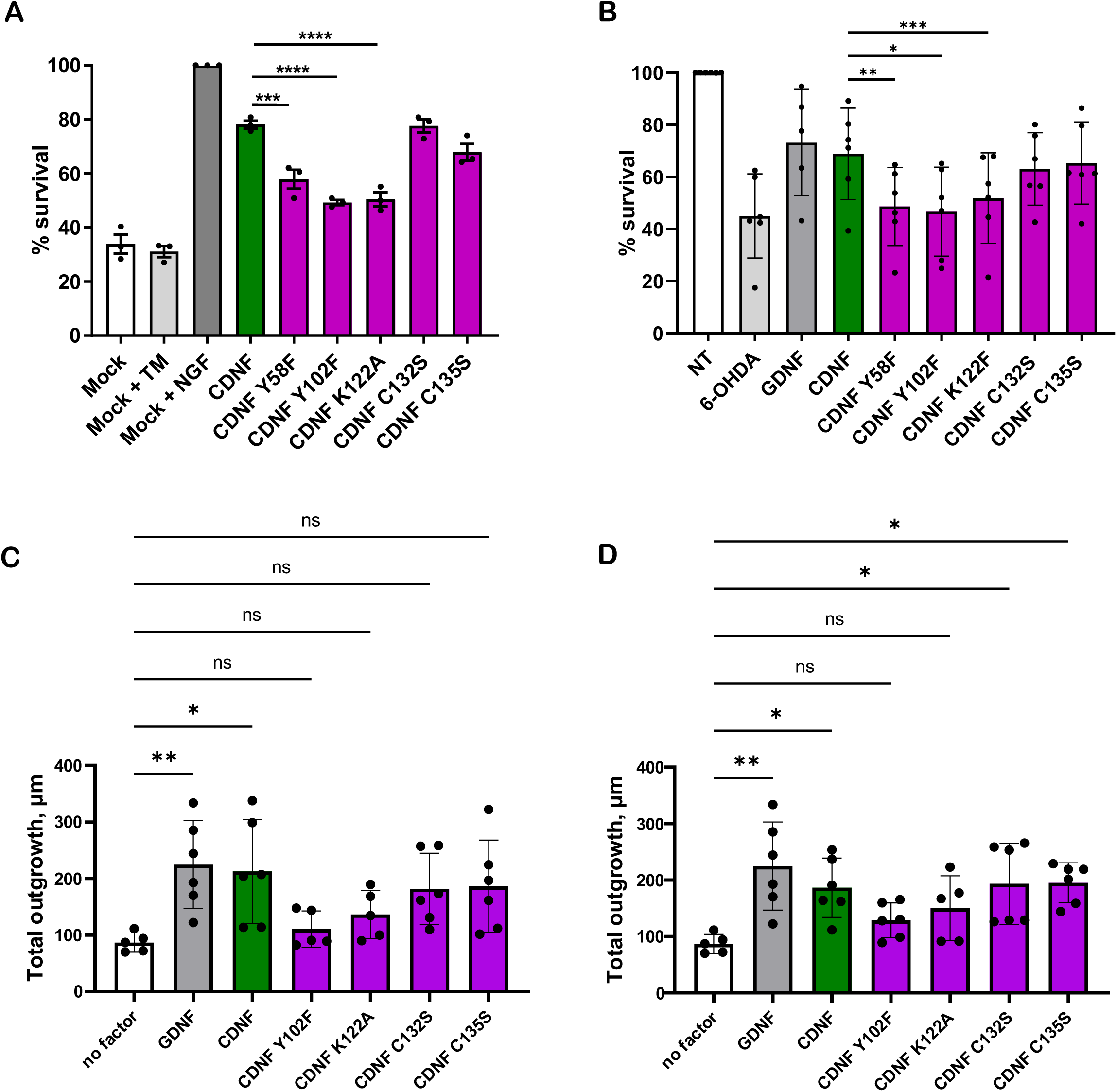
CDNF regulates neuronal survival and axonal regeneration through direct interaction with UPR sensors. (A) Microinjections of CDNF Y58F, CDNF Y102F or CDNF K122A purified proteins to SCG neurons do not rescue them from TM-induced cell death compared with mock-injected. TM was used at 2 µM for 72 hours. Mean ± SEM; n = 3. (B) Survival of human iPSCs-derived DA neurons with CDNF (100 ng/ml) or CDNF mutants (all at 100 ng/ml) in the presence of 6-OHDA (50µM). Data are presented as mean ± SEM. *p<0.01, **p<0.01, ***p<0.001. (C) Total neurite outgrowth of human iCell DopaNeurons after five days of treatment with 10 ng/ml GDNF, wild-type CDNF, and CDNF mutants (C132S, C135S, Y102F, and K122A). Neurite outgrowth was assessed by measuring total neurite length (µm) per cell, as quantified by immunostaining with anti-MAP2 and DAPI and analyzed using CellProfiler software. Statistical significance compared to the untreated control (no factor) is indicated: *p<0.05. Error bars represent the mean ± SD. (D) Total neurite outgrowth of human iCell DopaNeurons following treatment with the same proteins as in panel C with a concentration of 100 ng/ml. Statistical significance between groups is indicated: *p<0.05, **p<0.01. Error bars represent the mean ± SD.

### CDNF rescues human iPSCs-derived dopamine neurons through PERK and IRE1α

The direct effect of CDNF on human iPSCs-derived neurons has never been studied. To ensure the preclinical relevance of our findings in the mouse neuronal model, we further tested the neuroprotective activity of CDNF mutant proteins in human iPSCs-derived DA neurons. We set the conditions for maintaining these neurons in the medium without growth factors and conditions for survival assay, where combined oxidative and ER stress was induced using 6-OHDA. We found that CDNF in a dose-dependent manner can protect human dopamine neurons from 6-OHDA-induced cell death (Lindholm *et al.*, submitted). We then asked whether CDNF binding to IRE1α, or PERK is required for its neuroprotective activity in human neurons. Therefore, we tested CDNF mutants that cannot bind to IRE1α and mutants deficient in PERK binding. We found that in iPSCs-derived human DA neurons, both IRE1α binding deficient CDNF Y58F, CDNF Y102F, and PERK binding deficient CDNF K122A were lacking pro-survival and neuroprotective activity upon 6-OHDA-induced stress and cell death (Figure 6B). Cysteine loop mutants CDNF C132S and CDNF C135S protected iPSCs-derived human DA neurons from 6-OHDA-induced death similarly to wt CDNF and GDNF (Figure 6B). Since cysteine loop mutants CDNF C132S and C135S do not bind to BiP but protect both SCG and human DA neurons we conclude that CDNF binding to BiP is not required for its neuroprotective and neurorestorative activity. The results obtained on human DA neurons are in line with our results in mouse SCG neurons.

### CDNF promotes axonal regeneration in human iPSCs-derived dopamine neurons through interaction with PERK and IRE1α

In addition to preventing the dopamine neurons from dying, a potentially useful neuroregenerative property of a drug for PD is to promote neurite outgrowth, axonal regeneration and synapse formation. We tested whether CDNF possesses neuroregenerative properties in human iPSCs-derived DA neurons. The neurotrophic factor GDNF, known to efficiently support the survival of DA neurons and promote their neurite regeneration was used as a positive control. We found that CDNF at 100 ng/ml and even at 10 ng/ml increased the total outgrowth as efficiently, as GDNF at 100 ng/ml (Figure 5C). Additionally, CDNF at 10 ng/ml and 100 ng/ml increased total neurite outgrowth (Figure 1D). CDNF Y102F and CDNF K122A, deficient for IRE1α and PERK binding respectively, showed a compromised ability to promote total outgrowth both at 10 ng/ml and at 100 ng/ml, as compared with CDNF and cysteine loop mutants CDNF C132S and CDNF C135S (Figure 6C and 6D). Cysteine loop mutants CDNF C132S and CDNF C135S did not promote total neurite outgrowth at 10 ng/ml, but increased it at 100 ng/ml, similar to CDNF and GDNF (Figure 6C and 6D). No differences in mean branches and mean processes were observed at 10 ng/ml and 100 ng/ml for CDNF, GDNF, as well as for CDNF Y102F, CDNF K122A, CDNF C132S and CDNF C135S (Figures S6A-S6D).

## DISCUSSION

CDNF passed phase 1 clinical trials for the treatment of PD (NCT03295786,^22^). Currently, a peptide analogue of CDNF is in a new phase 1 clinical trial for PD treatment (NCT06659562). However, the precise molecular mechanisms underlying the therapeutic effects of CDNF remain unknown. While some studies link CDNF to the UPR^26,28,33^, the receptors and signalling pathways involved are still unclear. Most of the CDNF resides inside the cells in the ER lumen, and its secretion occurs during ER stress^48,49^. Additionally, ER stress triggers CDNF expression in cultured neurons^24^ and mouse liver and kidney tissue^26^. *In vitro* studies suggest that CDNF expression induces the adaptive UPR and suppresses apoptotic pathways^25^. Both intracellular and extracellular CDNF modulated UPR signalling and protected neurons from ER stress-induced cell death, induced by tunicamycin, thapsigargin, or 6-OHDA^26,31–33^.

A recent study where the mutagenesis analysis was performed with a C-terminal part of CDNF (61 amino acids of 161 amino acid protein) claims that the neuroprotective activity of CDNF relies on BiP binding without addressing whether the CDNF mutants that are putatively deficient for BiP binding interact with UPR sensors^41^. Our competition experiments demonstrate that the binding sites of BiP and CDNF in UPR sensors are overlapping. Therefore, the conclusions on the mechanism of neuroprotective biological activity could be made only if the corresponding mutants are analysed for binding to both BiP and UPR sensors. Notably, inhibiting IRE1α and PERK pathways blocked the neuroprotective action of CDNF in sympathetic and DA neurons^26^, but not the neuroprotective effects of BDNF and ciliary neurotrophic factor (CNTF)^33^. Similarly, IRE1α and PERK activities were demonstrated to be essential for the neuroprotective effect of CDNF in motor neurons^33^. Importantly, CDNF rescued motor neurons in three different animal models of ALS by suppressing the downstream signalling through all three UPR pathways, IRE1α, PERK and ATF6^33^. These *in vitro* and *in vivo* data suggest that the neuroprotective activity of CDNF occurs through the UPR sensors. These findings are in line with our current study, where through binding experiments, mutagenesis and neuroprotective assays, we demonstrated the importance of the direct binding of CDNF to PERK and IRE1α for its neuroprotective and regenerative activity in mouse and human neurons.

The pro-survival action of CDNF was not limited to neurons, CDNF was shown to enhance cardiomyocyte survival under TG-induced ER stress in cellular and rodent models and protect the heart from ischemia-reperfusion injury *in vivo*^50^. Proteomic analysis of protein-protein interactions confirmed the interactions of CDNF with ER luminal proteins like chaperons BiP and GRP170 and protein disulfide-isomerases PDIA1 and PDIA6^26^. These chaperones were shown to directly interact with UPR sensors and regulate their activity upon ER stress, demonstrating that UPR sensors are the main signalling hubs and decision-makers^51^. Multiple UPR regulators modulate the UPR in diverse ways. PDIA6 functions as an attenuator of both the IRE1α and PERK, through direct molecular interactions with cysteines in IRE1α and PERK^52^. Hsp47 directly binds to IRE1α but acts as an activator, modulating the UPR response through the displacement of BiP^53^.

Here we identify CDNF as a high-affinity, direct-binder to all three UPR sensors (PERK, ATF6, and IRE1α). Its competition with BiP for binding to UPR sensors shows the importance of CDNF as an ER stress attenuator. BiP plays a critical role in the initial stages of the UPR by binding to and inhibiting the UPR sensors PERK, IRE1α, and ATF6 under non-stressed conditions^54–56^. The dissociation of BiP from these sensors is a key step of UPR activation. Our data on CDNF-BiP competition suggest that CDNF binds to the same or overlapping binding site in UPR sensors as BiP and modulates UPR dynamics, similarly to BiP, through the attenuation of UPR response.

Competition of CDNF and MANF for IRE1α LD binding, suggests that CDNF and MANF share overlapping binding sites and may act similarly in binding and regulation of the activity of UPR sensors. However, since MANF concentrations in most tissues are significantly higher than that of CDNF, MANF binding to the UPR sensors may represent a main mechanism, and CDNF binding can complement it in chronic stress. In some tissues, such as skeletal muscles, where CDNF concentration is high, CDNF binding could determine the progression or slowing down of the pathology, related to hyperactivated UPR. In line with that, CDNF knockout resulted in the loss of enteric neurons^57^ and increased activation of UPR in the mouse muscle tissue, which in muscle was aggravated when MANF was ablated^32^.

As mentioned, the expression levels of CDNF in most tissues are about 10-100 times lower than those of MANF, complicating the study of endogenous CDNF protein. Attempts to perform co-immunoprecipitation (co-IP) were hindered by the low endogenous levels of CDNF in cells, rendering it undetectable with available antibodies. Using BiFC, we found that CDNF-PERK interaction in the cells had a stronger fluorescent signal as compared with that for IRE1α suggesting preferential binding of CDNF to PERK over IRE1α, however, this finding can result from higher PERK expression, as compared to that of IRE1α. Preference for PERK over IRE1α, however, is also supported by the finding that purified CDNF has the highest affinity to PERK in MST assays. A preferential interaction with PERK could mean that CDNF plays a role in chronic ER stress response, affecting the phosphorylation of protein synthesis initiator factor eIF2α and the downstream activation of ATF4, critical for cell survival and adaptation to stress. In contrast to our data on MANF-mediated signalling, we did not observe a significant impact of CDNF on the downstream genes regulated by the IRE1α pathway. Therefore,

MANF may have priority in binding to IRE1α over other UPR sensors, and CDNF may be the primary regulator of PERK, activated later when the ER stress is chronic and the MANF levels are decreasing. Using computational modelling, followed by the generation of CDNF mutants and their testing for UPR sensors and BiP binding and neuronal survival and neuroprotective assays, we identified critical binding sites and amino acids of CDNF responsible for PERK, IRE1α, and ATF6 binding. Using MST, we found that CDNF K122A mutant did not bind PERK. Since the UPR sensors are ER transmembrane proteins, to assess whether the mutants deficient for binding to the luminal domains of UPR sensors are deficient for binding to full-length UPR sensors, we tested their binding using BiFC in cells followed by flow cytometry-based quantification of the fluorescence intensity. Keeping more physiological setup in BiFC could favour the accessibility of binding sites, and the presence of other interacting partners, post-translational modifications, or localization within specific ER subdomains in the BiFC assay could influence binding and explain the difference with the MST data.

Based on the BiFC, representing the direct interaction in cells with the UPR sensors within their native cellular environment, we found that CDNF Y102F was deficient for IRE1α binding, this mutant is similar to earlier characterized MANF K96A mutant, deficient for IRE1α binding and located in the linker region^34^. Interestingly, this mutant retained the ability to bind to other UPR sensors and BiP, and therefore the specific contribution of IRE1α binding in the CDNF biological activity could be further addressed. CDNF K122A did not bind PERK, had slightly compromised ATF6 binding, and was deficient in BiP binding in cells, but not in MST. The crystal structure of MANF-BiP NBD complex reveals K117 (K138) in the interface of their interaction, supporting the functional importance of this motif, and likely also K122 in CDNF, in BiP binding^59^. CDNF C135S was binding IRE1α and PERK and its binding to ATF6 and BiP was compromised. Since the neuroprotective and neuroregenerative activity of CDNF C135S was not compromised, we conclude that the binding of CDNF to ATF6 and BiP is not crucial for its survival-promoting activity. Thus, this unique subset of mutants allowed us to distinguish the contribution of specific UPR branches and BiP to the neuroprotective activity of CDNF.

The biological activity studies in neurons revealed that mutations Y102F and K122A, located in the linker region and C-terminal domain, respectively, render CDNF biologically inactive in neuroprotection assays on mouse sympathetic and human dopamine neurons *in vitro*. Since CDNF Y102F was deficient for IRE1α LD binding, and CDNF K122A was deficient for PERK binding, we conclude that the biological activity of CDNF relies on binding to PERK and IRE1α.

These data are not exactly in line with a recent study, attributing the neuroprotective activity of CDNF to BiP (GRP 78) binding^41^. However, the mutagenesis in this study was conducted on C-CDNF peptides, and not on full-length proteins, making a direct comparison with the current study difficult. While the binding of CDNF mutants, potentially deficient in BiP binding, to UPR sensors was not examined, the downstream UPR signalling data strongly suggest that these interactions are compromised. Our data on the neuroprotective activity of CDNF K122A, based on BiFC-compromised not only in PERK and slightly in ATF6 but also in BiP binding do not exclude that CDNF-ATF6 and CDNF-BiP and binding may contribute to the neuroprotective activity of CDNF.

Reduced biological activity of CDNF Y58F, with N-terminal domain mutation, suggested that the N-terminus of CDNF could be also involved in interaction with UPR sensors. Therefore, using a full-length CDNF is crucial in any studies on the mode of action of this protein. Cysteine loop mutations in the C-terminal domain of CDNF specifically disrupted the interaction of CDNF with BiP and ATF6, but not with PERK or IRE1α. Since both CDNF cysteine mutants C132S and C135S retain neuroprotective activity, CDNF-BiP interaction is dispensable for the neuroprotective activity of CDNF.

We further addressed UPR receptors’ downstream pathways regulated by CDNF by measuring downstream targets of the UPR pathways in a multiple myeloma cell line, where the expression of CDNF is significantly higher than in other cell lines. In this cell line, CDNF attenuated *ATF4* expression, while the PERK-binding deficient K122A mutant did not affect it. Interestingly, both CDNF Y102F and CDNF K122A downregulated ATF4 similarly to wt CDNF under DTT, meaning reductive stress conditions, suggesting additional signalling mechanisms, independent of UPR sensors’ binding may contribute to the action of CDNF, depending on the stress conditions. This observation aligns with a recent study demonstrating the context-dependent activity of MANF^60^. MANF knockdown in cardiomyocytes increased cell death from DTT-mediated reductive ER stress, but not from non-reductive ER stresses caused by TG-mediated ER Ca^2+^ depletion or TM-mediated inhibition of ER protein glycosylation. Similar to MANF, the function of CDNF in the ER stress response may depend on the type of stress. Further investigation into the distinct pathways utilized by CDNF in different stress conditions, along with exploring potential interaction partners, is crucial to fully elucidate the mode of action of CDNF in different pathological conditions. However, these studies are hampered due to the low expression of CDNF in cell lines and tissues.

Interestingly, CDNF downregulated *ATF6* mRNA levels in DTT and TG-mediated ER stress but did not in TM-treated cells where CDNF K122A resulted in upregulation of ATF6 mRNA levels. This specific type of ER stress (e.g., protein N-linked glycosylation inhibition by TM) might influence the interaction of CDNF with the ER stress response machinery, leading to differential effects on ATF6. The mutation K122A might be crucial for the interaction of CDNF with ATF6 LD, leading to the observed upregulation of ATF6 under TM-induced stress. The finding that CDNF might influence ATF6 through PERK and IRE1α and vice versa aligns with multiple studies, confirming the crosstalk between the UPR sensors and UPR pathways. In several recent studies, various modes of crosstalk between UPR sensors were reported. PERK pathway was shown to attenuate the IRE1α pathway activity, resulting in apoptosis^61^, PERK through ATF4 was reported to enhance IRE1α expression. IRE1α via XBP1s signalling helps to sustain PERK expression during prolonged ER stress^62^, and PERK is a direct target of sXBP1^63^. ATF6, in turn, regulates the levels of XBP1, downstream to IRE1α^64^ and CHOP, downstream to PERK^65^. These findings show that UPR sensors could not be considered, as distinct parallel mechanisms but represent an interconnected system with mutual fine-tuning mechanisms. There could be additional pathways by which CDNF regulates ATF6 mRNA levels, independent of PERK and IRE1α interaction. The data suggests that CDNF regulates ATF6 differently depending on the type of stress.

CDNF was shown to regulate the levels of pro-inflammatory cytokines and decrease inflammation^20,28^. We tested whether CDNF regulates inflammation through UPR sensors upon TM-induced ER stress and found that CDNF upregulated the mRNA levels of inflammation-related genes: CCAAT/enhancer binding protein (CEBP), mediating the levels of anti-inflammatory and pro-inflammatory genes during the tissue repair; anti-inflammatory interleukin-4 (IL-4); *c-JUN*, regulating the activation of macrophages. We demonstrated that wt CDNF downregulates the transcript level of pro-inflammatory cytokine IL-4 (Figure 5H). This effect was mediated by IRE1α and PERK binding, as CDNF K122A, deficient for PERK binding and CDNF Y102F, deficient for IRE1α binding, lacked this activity. Given that chronic inflammation and ER stress are typical of many neurodegenerative conditions, the capacity of CDNF to influence both pathways represents a dual-targeted approach to modify the disease pathology.

In the survival and neuroprotective assays in SCG neurons and human iPSCs-derived DA neurons, CDNF K122A and linker mutant CDNF Y102F reduced the neuroprotective and survival-promoting activity of CDNF, while the N-terminal domain mutant exhibited a partial effect and cysteine loop mutants did not affect the pro-survival and neuroprotective activity of CDNF. Importantly, these findings were consistent across two neuron types and two species, suggesting that UPR regulation by CDNF may be universal for different cell types and species. This highlights the critical roles of the C-terminal and linker regions in CDNF’s ability to promote neuronal survival.

Our *in vitro* studies using human iPSCs-derived DA and mouse superior cervical ganglion sympathetic neurons exposed to ER stress inducers TM and 6-OHDA established CDNF as an ER-localized protein regulating UPR signalling and protecting neurons from cell death through the direct interaction with UPR sensor proteins, PERK and IRE1α. We employed microinjection of SCG neurons to deliver CDNF expression plasmids or purified proteins intracellularly. In this system, both wt CDNF and its cysteine loop mutants CDNF C135S, and CDNF C132S promoted the survival of SCG neurons treated with TM and similarly in human iPSCs-derived DA neurons treated with 6-OHDA.

Both CDNF and MANF have a C-terminal CXXC motif, commonly found in redox enzyme catalytic centers^66^. Notably, mutating this motif (CDNF C132S, CDNF C135S) did not significantly impact CDNF’s ability to counteract neuronal death induced by TM or 6-OHDA. Earlier we demonstrated that cysteine loop mutation in MANF did not affect its pro-survival and neuroprotective activity, suggesting a potentially similar mechanism of action between CDNF and MANF in neuroprotection, independent of the integrity of the cysteine loops. However, in other studies where MANF activity was assessed for the ability to rescue the larval lethality, the CXXC motif was crucial for DmManf^67^. The observed difference in the functional significance of the CXXC motif in our studies compared to those on MANF in flies might be attributed to differences in experimental models and indicate that neuronal loss is not crucial for fly larval lethality. While our findings suggest the CXXC is not necessary for CDNF’s neuroprotective activity in TM and 6-OHDA treated neurons, further research is needed to elucidate the specific functions of this motif.

Both CDNF and MANF contain (N-terminal) saposin-like domains. Since saposin-like proteins usually interact with lipids or membranes, the N-terminal domain likely mediates the CDNF and MANF interaction with lipids^36,68^. Indeed, MANF was shown to directly bind sulfatide possibly via its N-terminal domain^69^. Interestingly, mutation within the N-terminal domain partially compromised CDNF neuroprotective activity, indicating that N-CDNF interaction with IRE1α may be involved in the pro-survival activity of CDNF.

In this study, we showed for the first time that CDNF has direct effects on human neurons promoting neurite outgrowth and survival of human iPSCs-derived DA neurons. We found that not only the neuroprotective but also the regenerative action of CDNF occurs through its interaction with UPR sensors. This finding is important for the clinical translation of CDNF and its analogues, particularly in the context of PD, a condition characterized by the degeneration and loss of dopamine neurons.

The ability of CDNF to induce neurite outgrowth suggests a direct role in supporting neuronal health and connectivity, which is crucial for the recovery and maintenance of neuronal function in neurodegenerative diseases. Given the progressive nature of PD and the lack of treatments that can halt or reverse neuronal degeneration, the implications of our findings are profound.

These properties position CDNF and its analogues, as very attractive drug candidates for PD. Unlike current therapies primarily focusing on symptom management, CDNF offers a therapeutic avenue aimed at the underlying causes, neuronal loss, neurite degeneration and dysfunction. As it regulates UPR, reduces inflammation, promotes regeneration and acts only on stressed cells it has rather unique properties among putative novel drugs for neurodegenerative diseases. Since CDNF protein has limited pharmacokinetic properties, the development of small molecule analogues that mimic CDNF activity broadens the therapeutic potential, offering a more versatile and potentially accessible treatment option for PD.

## LIMITATIONS OF THE STUDY

Due to a low level of expression of endogenous CDNF in cell lines and tissues, the current study lacks data on the interactions between endogenous CDNF and UPR sensors. We have provided a demonstration of the direct interactions, using multiple techniques to ensure the reproducibility of the results regardless of the used approach. The study would have benefited from testing the effects of CDNF and CDNF mutants in IRE1α and PERK knockouts, preferably in DA and SCG neurons. These will be the focus of a follow-up study. Mouse knock-in models of CDNF Y102F and CDNF K122A will bring more light on the importance of interaction with each of the UPR sensors at the organismal level.

## ACKNOWLEDGEMENTS

The study was supported by grants from the Finnish Research Council (Grant No 343299), and Sigrid Jusélius Foundation to MS, and from Jane and Aatos Erkko Foundation (to TK and MS). We thank Dr. Päivi Lindholm, Dr. Maria Lindahl, and Dr. Brandon Harvey for their critical comments on the manuscript. Support from the Swedish National Infrastructure for Biological Mass Spectrometry (BioMS) and the SciLifeLab, Integrated Structural Biology (ISB) platform, is gratefully acknowledged.

## AUTHOR CONTRIBUTIONS

V.K. and O.S. designed and performed MST, BiFC, WB, and RT-qPCR experiments. L.Y. performed neuronal survival and neurite outgrowth studies in SCG and human iPSCs-derived DA neurons. L.I. and M.K. planned and performed protein-protein docking experiments and wrote the respective part of the manuscript. S.F., L.H. and T.K designed and performed the crosslinking-MS experiments, and S.F and TK performed MS-based molecular modelling of the PERK-CDNF complex and wrote the respective part of the manuscript. S.F. purified BiP NBD for MST studies and PERK LD for MS crosslinking studies. U.T. and M.U. purified CDNF and IRE1α LD mutant and wild-type proteins. M.S. initiated the study, supervised the experiments and participated in the designs and manuscript writing. MS provided funding. All the authors read and approved the final version of the manuscript.

## DECLARATION OF INTERESTS

The authors declare no competing interests. MS is the inventor of the CDNF-related patents that belong to Herantis Pharma Plc. MS is also a shareholder in Herantis Pharma Plc.

## SUPPLEMENTARY FIGURES

**Figure S1. Expression and secretion of putatively deficient for UPR sensors binding CDNF mutant constructs in CHO cells.**

(A) SDS-PAGE gel electrophoresis of purified from CHO cells human recombinant CDNF and CDNF mutant proteins (bands 1 and 2-9, correspondently), non-reducing conditions, Coomassie blue staining. Molecular weight markers in kDa on the left.

(B) Quantification of the expression level of human recombinant CDNF and CDNF mutant proteins, containing HA-tags. HA-tag level was normalized to α-tubulin level, n = 3. One-way ANOVA, Tukey’s post hoc test. ∗p ≤ 0.05; ns, not significant.

**Figure S2. Proximity ligation assay (PLA) controls for CDNF interactions in HEK293T cells.**

(A) Proximity ligation assay (PLA) controls for CDNF interactions with IRE1α in HEK293T cells.The graph shows the number of dots per cell for various conditions: no antibody (No Ab) GFP, GFP-IRE1α with and without doxycycline (Dox), and CDNF-IRE1α without Dox. Mean dots/cell values ± SEM from n = 3 independent experiments are indicated. Statistical analysis was performed using Student’s t-test: **: p < 0.01.

(B) Proximity ligation assay (PLA) controls for CDNF interactions with PERK in HEK293T cells. The number of dots per cell under different conditions: no Ab GFP, GFP-PERK with and without Dox, and CDNF-PERK without Dox. Controls included antibody conditions without antibodies and the effect of Dox. Mean dots/cell values ± SEM from n = 3 independent experiments are indicated. Statistical analysis was performed using Student’s t-test: **: p < 0.01

(C) BiFC controls for signal formation in the nucleus and ER. The positive controls: transcription factors Jun and Fos for nuclear signal detection and the ER proteins GRP78 and PERK for ER-localized signals. Non-interacting transcription factors Jun and Max were used as negative controls for BiFC signal formation.

(D) Estimation of the expression level of Venus constructs of IRE1α, ATF6, and PERK/PERK KDD for bimolecular fluorescence complementation (BiFC) assay in HEK293 cells using western blotting (WB). A representative image is on the upper panel and quantification is on the lower panel. Venus level was normalized to α-tubulin level, n = 3. One-way ANOVA, Tukey’s post hoc test. ∗p ≤ 0.05; ns, not significant.

(E) Estimation of the expression level of Venus constructs of CDNF, N-CDNF, and C-CDNF for bimolecular fluorescence complementation (BiFC) assay in HEK293 cells using WB. A representative image is on the right panel and quantification is on the left panel. Venus level was normalized to α-tubulin level, n = 3. One-way ANOVA, Tukey’s post hoc test. ∗p ≤ 0.05; ns, not significant.

(F) Estimation of the expression level of Venus constructs of CDNF, CDNF Y58F, CDNF K115A, CDNF C132S, CDNF C135S, CDNF Y102F, and CDNF K122A for bimolecular fluorescence complementation (BiFC) assay in HEK293 cells using WB. A representative image is on the right panel and quantification is on the left panel. Venus level was normalized to α-tubulin level, n = 3. One-way ANOVA, Tukey’s post hoc test. ∗p ≤ 0.05; ns, not significant.

**Figure S3. Effect of CDNF and CDNF K122A overexpression and siRNA knockdown on IRE1α and eIF2α phosphorylation following tunicamycin treatment.**

(A, B, C, D) Western blot analysis was conducted to evaluate the phosphorylation status of IRE1α and eIF2α after tunicamycin treatment. Overexpression of CDNF and the CDNF K122A mutant (A, B), as well as siRNA knockdown of CDNF or CDNF/MANF (C, D), do not change in the phosphorylation levels of IRE1α and eIF2α, n = 3. One-way ANOVA, Tukey’s post hoc test. ∗p ≤ 0.05; ns, not significant.

**Figure S4. Analysis of cross-linking in the CDNF-PERK complex**

(A) Cross-linked residues used in docking; cross links are shown as yellow lines between labeled residues (distance range from 7 – 22 Å).

(B) Crosslinking of the CDNF-PERK complex. SDS-PAGE analysis of crosslinked CDNF-PERK complexes. Lane 1 (first well): No crosslinker added; lane 2: 0.5 mM DSS crosslinker added; lane 3:1 mM DSS crosslinker added.

**Figure S5. qPCR analysis of unfolded protein response genes following CDNF and CDNF mutant treatment.**

Treatment of HEK293 cells with CDNF or the CDNF K122A mutant did not change the mRNA levels of stress response genes *CHOP, GRP78*, and *sXBP1* after tunicamycin (TM) treatment (A-C), DL-Dithiothreitol (DTT) (D-F) or thapsigargin (TG) (G-I) treatment. Cells were treated for 4 hours with TM (2.5 µg/ml), TG (1 µM), or DTT (2 mM), followed by 20 hours of treatment with 400 nM CDNF, CDNF K122A, or CDNF Y102F. Data are presented as mean ± SEM from 3 to 4 independent experiments, with statistical analysis performed using one-way ANOVA followed by Tukey’s post hoc test.

**Figure S6. Human DA neuron neurite outgrowth assay**

(A) mean processes and (B) mean branches per cell following the treatment with CDNF (10 ng/ml) or CDNF mutants (10 ng/ml) and (C) mean processes and (D) mean branches per cell following the treatment with CDNF (100 ng/ml) or CDNF mutants (100 ng/ml). Data are presented as mean ± SEM. *p < 0.05, ** p < 0.01, *** p < 0.001, **** p < 0.0001.

## RESOURCE AVAILABILITY

## EXPERIMENTAL MODEL AND SUBJECT DETAILS

### Cell lines

HEK293 cells for bimolecular fluorescence complementation assay (BiFC) experiments were cultured in Dulbecco’s modified Eagle’s medium (DMEM). HEK293 and U2OS cells used for plasmid DNA or siRNA transfection experiments were grown in Minimum Essential Media (MEM 61100-087, Gibco). Flp-In T-REx 293 cell line (Invitrogen) containing a single stably integrated FRT site and expressing Tet repressor were used for the generation of inducible cell lines, expressing IRE1α-HA and GFP-HA. The medium composition was the same as for HEK293 cells. CHO cells were grown in growth media consisting of Ham’s F12 nutrient mix (21765029, Thermo Fisher Scientific), 2 mM GlutaMAX (35050061, Thermo Fisher Scientific). CHO cells were used to analyse the secretion and expression of CDNF mutant proteins. RPMI8226 cells (kindly provided by Dr. Caroline Heckman) were cultured in RPMI-1640 (ThermoFisher Scientific, Massachusetts, United States) at 37 °C and 5% CO_2_. HeLa cells were used for proliferation assay, were cultured in in Dulbecco’s modified Eagle’s medium (DMEM), All the cell lines used were maintained in media supplemented with 10% FBS (10270106, Gibco) and 50 mg/ml normocin (ant-nr-2, InvivoGen) at 37 ^0^C and 5% CO_2_ incubator.

### Reagents and proteins

The endoplasmic reticulum (ER) stress-inducing agents, tunicamycin (TM) (ab120296, Abcam) and thapsigargin (TG) (T7459, ThermoFisher Scientific) and inhibitor of protein synthesis cycloheximide (C1988-5G, Sigma) were prepared by dissolving them in dimethyl sulfoxide (DMSO, Sigma) and were stored at a temperature of −20°C. These agents were subsequently diluted in the cell culture medium, ensuring the final concentration of DMSO was less than 0.1%, immediately before their application.For survival assays in neurons neurotoxin 6-hydroxydopamine hydrochloride (6-OHDA, H4381, Sigma Aldrich) was used. For the IRE1α oligomerization assay, we used two inhibitorsIRE1α: KIRA6 (HY-19708, MedChemExpress) and 4μ8C (14003-96-4, Cayman Chemical). LDs of three UPR sensors, human IRE1α, PERK, ATF6 and IRE1α LD mutant proteins were expressed and purified in CHO cells by Icosagen Ltd. (Tartu, Estonia) and described in (Kovaleva et al. 2023). Human recombinant GRP78 (BiP) was obtained from StressMarq Biosciences Inc. (SMB-SPR-119A). Human recombinant BiP NBD and PERK residues 95-420 (PERK_95-420_) were purified from *E. coli*. Recombinant human CDNF protein and human CDNF mutant proteins were expressed and purified from a CHO-derived cell line Icosagen Ltd., Tartu, Estonia. Some of the CDNF used in this study was purified from CHO cells using a similar approach by Biovian, Turku, Finland

## METHOD DETAILS

### Microscale thermophoresis (MST)

The study utilized the Monolith NT.115 instrument (NanoTemper Technologies GmbH, Germany) to evaluate the binding affinities of recombinant proteins at 25°C. The proteins for MST were purified from CHO cells, and α-synuclein from *E. coli*, The CDNF and PERK proteins were labeled using His-Tag Labeling Kit RED-tris-NTA (MO-L008) and the amine-reactive Monolith Protein Labeling Kit RED-NHS 2nd Generation (L011), respectively. The unbound dye was removed post-labeling using Zeba Spin Desalting Columns (89882, Thermo Fisher Scientific), following manufacturer protocols. All proteins were assessed at 20 nM across experiments, in an MST buffer composed of 10 mM Na-phosphate (pH 7.4), 1 mM MgCl_2_, 3 mM KCl, 150 mM NaCl, and 0.05% Tween-20. The MST measurements leveraged premium coated capillaries (NanoTemper Technologies GmbH) and optimized 100% LED power settings for detection. Data collection spanned 12-14 data points for each ligand concentration curve, with each point averaging results from 3-5 independent assays to calculate normalized fluorescence changes (ΔFnorm), standard deviations, and dissociation constants (Kd) with error margins. Analysis was conducted using MO. Affinity Analysis software (versions 2.3 and 2.2.4) and GraphPad Prism (versions 8.0.2 and 9) s.

### Crosslinking of the CDNF-PERK_95-420_ complex

For crosslinking, 105 μM (5.3 μg) of CDNF was mixed with 105 μM (10.6 μg) of PERK_95-420_ in a final reaction volume of 5.6 μl in 1xPBS buffer complemented with 2 mM MgCl_2_, and incubated for 10 min, 22°C, 500 rpm in a thermoblock for the proteins to bind to each other. Disuccinimidyl suberate (DSS-H12/DSS-D12, Creative Molecules Inc., 001S) was added to final concentrations of 0.5 or 1.0 mM sequentially (0.25 mM + 0.25 mM or 0.5 mM + 0.5 mM, respectively) and the samples further incubated for 30 min, 22°C, 500 rpm after each addition. The crosslinking reaction was quenched with a final concentration of 50 mM of ammonium bicarbonate for 15 min, 22°C, 500 rpm. Samples were prepared both for in-solution digestion and for extraction of crosslinked CDNF-PERK_95-420_ complexes from SDS-PAGE gels.

### Preparation of crosslinked samples for mass spectrometry analysis

In-solution digested samples for MS analysis were prepared essentially as described^71^. The proteins were denatured using 8 M urea – 100 mM ammonium bicarbonate, and the cysteine bonds were reduced with a final concentration of 5 mM Tris(2-carboxyethyl) phosphine hydrochloride (TCEP, Sigma, 646547) for 60 min, 37°C, 800 rpm. Denatured and reduced samples were subsequently alkylated using a final concentration 10 mM 2-iodoacetamide for 30 min, 22°C in the dark. For digestion, 1 μg of lysyl endopeptidase (LysC, Wako Chemicals, 125-05061) was added, and the samples were incubated for 2 h, 37°C, 800 rpm. The samples were diluted with 100 mM ammonium bicarbonate to a final urea concentration of 1.5 M, and 1 μg of sequencing grade trypsin (Promega, V5111) was added for 20 h at 37°C, 800 rpm. The digested samples were acidified with 10% formic acid to a final pH of 3.0. Peptides were purified and desalted using C18 reverse phase column following the manufacturer’s recommendations (The Nest Group, Inc.). Dried peptides were reconstituted in 20 μl of 2% acetonitrile and 0.1% formic acid before MS analysis. Alternatively, the crosslinked samples were separated using 4-20% SDS-PAGE (Bio-Rad, Mini-PROTEAN TGX Precast Protein Gels, 4561096), and the protein bands corresponding to the 1:1 ratio CDNF-PERK_95-420_ complexes excised and prepared for MS as described^72^. The extracted peptides were reconstituted in 10 μl of 2% acetonitrile and 0.1% formic acid before MS data acquisition.

### Liquid chromatography-mass spectrometry for crosslink identification

For all samples, 1 μl of peptides were analyzed on an Orbitrap Eclipse mass spectrometer connected to an ultra-high-performance liquid chromatography Dionex Ultra300 system (Thermo Scientific). The peptides were loaded and concentrated on an Acclaim PepMap 100 C18 precolumn (75 μm × 2 cm) and then separated on an Acclaim PepMap RSLC column (75 μm × 25 cm, nanoViper, C18, 2 μm, 100 Å; both columns Thermo Scientific), at a column temperature of 45°C and a maximum pressure of 900 bar.

A linear gradient of 5–25% of 80% acetonitrile in aqueous 0.1% formic acid was run for 100 min followed by a linear gradient of 25–45% of 80% acetonitrile in aqueous 0.1% formic acid for 20 min. One full MS scan (resolution 120,000; mass range of 400–1,800 *m/z*) was followed by MS/MS scans (resolution 30,000) with a 3 sec cycle time. Precursors with an unknown charge state, a charge state of 1, 2, or above 9 were excluded. The precursor ions were isolated with 1.6 *m/z* isolation window and fragmented using stepped higher-energy collisional-induced dissociation (HCD) at a normalized collision energy of 21, 26, 31. The dynamic exclusion was set to 60 s.

### Crosslinking data analysis

All spectra from crosslinked samples were analyzed using pLink 2 (version 2.3.11)^73^. The target protein database contained the sequence for the CDNF-PERK_95-420_ proteins including common contaminants. pLink2 was run using default settings for conventional HCD DSS-H12/D12 crosslinking, with trypsin as the protease and up to three missed cleavages allowed. Peptides were selected with a mass between 600 and 6,000 Da, and a length between 6 and 60 amino acids. Precursor and fragment tolerance were set to 20 and 20 ppm, respectively. Crosslink identifications were filtered by requiring 10 ppm mass accuracy, false discover rate (FDR) < 5%, and an *E*-value < 0.01. The data have been deposited to the ProteomeXchange consortium via the MassIVE partner repository (https://massive.ucsd.edu/) with the dataset identifier PXD057345.

### Generation of CDNF mutant plasmids and recombinant proteins

pcDNA5/FRT/TO pre-SH-CDNF Y58F, Y102F, K115A, C132S, C135S and K122A mutants were generated using site-directed inverse PCR mutagenesis and pcDNA5/FRT/TO pre-SH-CDNF as template. The pcDNA5/FRT/TO pre-SH-CDNF Y102FK122A and K115AK122A mutants were generated using pcDNA5/FRT/TO pre-SH-CDNF K122A as a template. The wt CDNF and CDNF Y58F, Y102F, C132S, C135S and K122A mutant recombinant proteins were produced by Icosagen (https://www.icosagen.com/) using a mammalian protein expression system from a CHO-derived cell line, as has been described before^74^. Some of the CDNF used in this study was produce dby Biovian, Turku, Finland using similar technology. Briefly, codon-optimized cDNAs were cloned to pQMCF-T expression vectors which were then transiently transfected to CHO-derived protein production cell line. Proteins were captured and purified from the cell culture media using 5 ml Q FF followed by 1 ml SP HP, buffer was exchanged into PBS, pH 7.4, by size exclusion chromatography. Protein purity was verified by SDS-PAGE with Coomassie staining and immunoblotting using polyclonal rabbit anti-CDNF antibody (300-100, Icosagen).

### Expression and secretion of CDNF mutant plasmids

CHO cells were grown on 6 cm plates and transfected with pTO expression plasmids 6 µg using PEI transfection reagent 1 µg/µl in 1× PBS, pH 4.5; 4:1 v/w ratio of PEI:DNA (PEI, Polysciences) per plate. Media was changed 24 hours after transfection to serum-fee media (3 ml per 6 cm plate). Incubated for 24 hours more before harvesting cells and collecting media. Each cell pellet was lysed in 400 µl of lysis buffer, 100 µl of media was set aside before concentrating, and the rest was concentrated from ∼ 3 ml to ∼ 100 µl using Amicon Ultra-4 centrifugal filters 10K. 20 µl of each sample was loaded onto 4 - 15% gel. Primary antibody: anti-HA (Abcam) 1:1000, 2 hours. Secondary antibody: goat-anti-mouse 690 LR (Licor) 1:10000, 24 hours.

### Duolink proximity ligation assay (PLA)

The experiments were performed on a 96-well format on Flp-In T-REx 293 cells, expressing CDNF-HA/GFP-HA upon doxycycline induction. 1 × 10^4^ cells/well were plated on pre-coated with poly-D-Lysine (0.1 mg/ml) black Perkin Elmer plates. Cells were fixed with 4% paraformaldehyde for 15 min and afterwards permeabilized/stained with DAPI (D9542, Sigma-Aldrich) in 1 x PBS containing 0.05% Triton X-100 for 10 min. Blocking and incubation with antibodies have been performed following Duolink manufacturer’s protocol. Cells were incubated overnight at 4°C with the following primary antibodies: anti-PERK rabbit pAb (CST, 3192), anti-IRE1α rabbit mAb (CST, 3294), anti-BiP rabbit mAb (CST, 3177), anti-HA mouse mAb (Abcam, ab130275). Incubation with PLUS and MINUS PLA probes have been performed for 1 hour at 37°C. Ligation and amplification were performed according to the manufacturer’s instructions. The imaging of 16 sites/well was performed in TexasRed and DAPI channels using MolecularDevices Nano scanner. The analysis and quantification were done using CellProfiler 3.1.5 and CellProfiler Analyst 2.2.1 software^75^.

### Plasmids for BiFC

The generation of pEZY BiFC Jun-NV, pEZY BiFC Max-CV, pEZY BiFC Fos-CV, pEZY BiFC Grp78-NV(CV), pEZY BiFC pre-CDNF-NV(CV), pEZY BiFC pre-NV-N-CDNF, pre-NV-C-CDNF and pEZY BiFC IRE1a-NV(CV) has been described in detail before^34,76^. CDNF mutants were generated using site-directed inverse PCR mutagenesis using pEZY BiFC CDNF-NV(CV) as a template. In a preliminary step of generating PERK constructs, the pEZY BiFC pre-CDNF-NV template was linearized, and the insert removed through inverse PCR using 5′-phosphorylated primers. This template was then treated with DpnI enzyme (Thermo Fisher) to digest it. The resulting product was separated on an agarose gel, excised, and purified using the NucleoSpin Gel and PCR Clean-Up Mini Kit (Macherey-Nagel, Düren, Germany). Following this, the purified product was ligated using T4 DNA ligase (Thermo Fisher) obtaining pEZY BiFC-stop destination vector. We created two versions of BiFC constructs for the PERK protein: one is the wild-type PERK (PERK wt) and the other is a modified PERK (PERK KDD), which contains a one-site mutation in the kinase domain rendering the kinase inactive. Both versions were linked to either the N-terminal (VN173) or C-terminal (VC155) segment of the Venus fluorescent protein. To assemble the pEZY BiFC pre-NV(CV)-PERK and pEZY BiFC pre-NV(CV)-PERK KDD constructs, we amplified the PERK/PERK cDNA from plasmids gifted from David Ron (Addgene #21814 and #21815). We also amplified the VC155 (C-Venus) and VN173 (N-Venus) sequences from the respective BiFC vectors. The sequences were inserted into the pEZY BiFC-stop vector between the regions encoding the signal peptide and the mature human PERK. This was accomplished using NEB-builder HiFi DNA assembly, which allowed to cloning of PERK, PERK KDD, and selected Venus fragments into the pEZY BiFC vector, resulting in four final constructs.

### Bimolecular fluorescence complementation assay (BiFC)

HEK293 cells were seeded on coverslips coated with poly-D-lysine (P0899, Sigma-Aldrich) and, after 48 hours, co-transfected with pEZY BiFC N-Venus and C-Venus plasmids using jetPEI transfection reagent (101, Polyplus-transfection) following the manufacturer’s instructions. After transfection, the cells were fixed using 4% paraformaldehyde (PFA), rinsed with PBS, and then permeabilized with 0.1% Triton X-100 in PBS (PBS-T). For staining nucleus and endoplasmic reticulum, we employed ER-ID Red assay kit (ENZ-51026-K500, Enzo Life Sciences), containing Hoechst 33342 for nuclear staining and ER-ID Red detection reagent. The coverslips were mounted using ProLong Diamond Antifade Mountant (P36965, Thermo Fisher Scientific). Imaging was conducted on a Leica SP8 STED confocal microscope using a 63x glycerol immersion objective and the Leica Application Suite X (LASX) software. Image processing, including uniform adjustments to brightness and contrast for all images, was performed with CorelDRAW 2018.

### Fluorescence-activated cell sorting (FACS)

HEK293 cells were seeded at a density of 0.1 to 0.2 x 10^6^ cells per well on a 12-well plate. The following day, the cells were transfected with BiFC CV and BiFC NV constructs of the proteins of interest. After 48 h transfection, TrypLE without phenol red (12604021, Thermo Fisher) was applied to each well. This was then neutralized using a media (Opti-MEM without phenol red + 10% FBS). Next, the cell suspension was centrifuged, the supernatant aspirated and the pellet reconstituted in PBS. The resulting cell suspension was then passed through cell strainer caps into polypropylene tubes (both supplied by BD Biosciences) to remove any clumps. Finally, the fluorescence of the samples was measured using a FACS Fortessa flow cytometer (BD Biosciences).

### Fluorescence measurement using flow cytometry

Fluorescence intensities of cell samples were measured using a LSRFortessa (BD) flow cytometer, equipped with a 100 µm nozzle and a 488 nm blue excitation laser. For each measurement, 500 µl of well-resuspended cell culture (OD600 = 0.5) was loaded into the flow cytometer. Prior to analysis, debris, and non-uniform cells were excluded through appropriate gating. Fluorescence intensities were recorded from 10,000 live cells using the GFP channel (527/32 nm bandpass filter) with excitation at 488 nm. To differentiate Venus signals from background autofluorescence, a threshold of 500 fluorescence intensity units was set. This ensured that only relevant signals were captured, providing a clear distinction between autofluorescence and specific Venus fluorescence.

### Protein-protein docking

The structure with the least restraint violations from the NMR solution structure of the recombinant, full-length of CDNF (PDB ID: 4BIT)^39^ and the crystal structure of the PERK LD (PDB ID: 4YZS)^40^ were used for docking study. The 3D structures of the CDNF – PERK LD complexes were predicted by protein-protein docking procedure using Schrödinger LLC BioLuminate software (Schrödinger Release 2018-3: BioLuminate, Schrödinger, LLC, New York, NY (2018). The 3D structures of the studied proteins were treated using the Protein Preparation Wizard (OPLS_2005 force field) in the Schrödinger LLC Maestro software (Schrödinger Release 2018-3: Schrödinger Suite 2018-3 Protein Preparation Wizard; Epik, Schrödinger, LLC, New York, NY (2018); Impact, Schrödinger, LLC, New York, NY, 2018; Prime, Schrödinger, LLC, New York, NY (2018). The protein-protein docking was carried out using the PIPER procedure implemented in Schrödinger LLC BioLuminate software (Schrödinger Release 2018-3: BioLuminate, Schrödinger, LLC, New York, NY (2018), which proceeds with a rigid body global search based on the Fast Fourier Transform (FFT) correlation approach^77^. The PIPER procedure performs exhaustive evaluation of an energy function in discretized 6D space of mutual orientations of two proteins. The structures corresponding to different mutual orientations of the proteins were ordered according to the scoring function that is given as the sum of terms representing shape complementarity, electrostatic, and desolvation contributions The top 1000 structures were subsequently clustered using the pairwise root mean square deviation (RMSD) as the distance measure between two proteins in the complex within a fixed clustering radius 9Å^78^. The selected structures in 30 largest clusters were refined by a Semi-Definite programming-based Underestimation medium-range optimization method^79^. The analysis of the protein–protein interactions in the predicted 30 CDNF – PREK LD complex configurations was performed using AutoDock Tools 1.5.6 software^80^.

### IRE1α oligomerization assay

TREx-293 IRE1α-3FGHGFP cells^35^ were plated 5 × 10^3^ cells/well on pre-coated with poly-D-lysine (0.1 mg/ml) black Perkin Elmer plates in DMEM with 10% FBS and 100 µg/ml normocin. Next day the cells were transiently transfected with pTO-pre-SH-CDNF-GW-FRT (MANF mutants) or pTO-SH-GW-FRT as a control vector, 100 ng of plasmid/well using PEI Transfection Reagent (1 µg/µl in 1x PBS pH 4.5; 4:1 v/w ratio of PEI:DNA). 24 hours after transfection, IRE1α-GFP expression was induced with doxycycline (1 µg/ml) treatment for 24 hours. ER stress was induced by treating the cells with the inhibitor of N-linked glycosylation tunicamycin (TM), 5 µg/ml for 4 hours. Thereafter, cells were fixed with 4% PFA for 20 min and stained with DAPI (D9542, Sigma-Aldrich) in 1 x PBS for 10 min. Imaging (16 sites/well) was performed using MolecularDevices Nano scanner. Three independent experiments were analyzed and quantified using CellProfiler 3.1.5 and CellProfiler Analyst 2.2.1 software^75^.

### RNA interference

For transient knockdown of CDNF or MANF in HEK293/HeLa cells, the cells were seeded at a density of 2.8 x10^5^ cells/well on 12-well plates. siRNAs targeting CDNF (siGENOME SMARTpool Human siRNA CDNF, 441549) and MANF (7873), both obtained from Horizon Discovery, were used to specifically decrease the expression of CDNF, MANF, or both proteins simultaneously. As a negative control, the siGENOME Non-Targeting siRNA Control Pool #1 (Dharmacon, Horizon Discovery) was utilized. The transfection of HEK293/HeLa cells with the selected siRNAs was performed using Dharma-FECT reagent 1, following the manufacturer’s instructions (Dharmacon, Horizon Discovery). Between 24 to 48 hours post-transfection, the cells were prepared for subsequent analyses.

### Transient transfections with plasmids

HEK293/U2OS cells were plated 2.5 × 10^5^ cells/well on 6-well plates in DMEM with 5% FBS. Next day the cells were transfected using PEI transfection reagent (24765-100, Polysciences) 1 mg/ml in 1x PBS, pH 4.5; 4:1 v/w ratio of PEI:DNA with plasmids carrying CDNF or CDNF mutant. After 24 hours of transfection cells with plasmid DNA cells were treated with TM 2.5 µg/ml for 2 hours, 4 hours, 6 hours, 20 hours, followed by transfection with plasmids carrying CDNF treatment with exogenous human recombinant CDNF (400 nM) for 20 hours.

After 48 hours after transfection cells were harvested and processed for further experiments.

### Western blot analysis

HEK293T cells were plated 2.5 × 10^5^ cells/well on 6-well plates in DMEM with 5% FBS. Next day the cells were transfected using PEI transfection reagent (24765-100, Polysciences) with plasmids carrying CDNF or CDNF mutant or with siRNA using Dharma-FECT reagent 1. After 24 hours of transfection cells with plasmid DNA and 48 hours after transfection with siRNA cells were treated with TM 2.5 µg/ml for 2 hours, 4 hours, 6 hours, 20 hours, followed by transfection with plasmids carrying CDNF treatment with exogenous human recombinant CDNF (400 nM) for 20 hours. The lysis was performed in RIPA buffer, containing protease and phosphatase inhibitor cocktail tablets (Roche). The concentrations of total protein in cell lysates were measured using NanoDrop 2000 spectrophotometer. 20 µg/well of total protein was loaded onto Bio-Rad mini-PROTEAN precast gels followed by the transfer onto nitrocellulose membrane. The transfer was performed for 1 hour on ice at RT. The membranes were blocked with 5% BSA TBS-T (or 5% milk TBS-T) for 1 hour at RT and then incubated with anti-IRE1α (CST, 3294), IRE1α pS724 (PA1-16927, Invitrogen), α-Tubulin (Sigma-Aldrich, T9026) primary antibodies overnight at 4°C. Peroxidase-coupled secondary antibodies and the enhanced chemiluminescence (ECL) detection system were used for western blot development.

### Real-time quantitative PCR (RT-qPCR)

RPMI8226 cells were plated 2.5 × 10^5^ cells/well on 6-well plates in RPMI 1640 with 5% FBS. Next day the cells were treated with TM 2.5 µg/ml for 4 hours, followed by treatment with exogenous human recombinant CDNF (400 nM), mutant CDNF K122A and CDNF Y102F (400 nM) for 20 hours. RNA was extracted from the RPMI8226 cells using a TRI reagent (AM9738, Invitrogen) with DNase I (EN0521, Thermo Fischer) treatment. TRI reagent was added to the samples, incubated for 5 min, collected and chloroform (1:5) was added. The samples were centrifuged at 13000 rpm for 15 min, the aqueous phase of the samples was collected and mixed with isopropanol (1:1) and incubated for 10 min. Thereafter, the samples were centrifuged at 13000 rpm for 10 min and washed with 75% ethanol. After the centrifugation at 8000 rpm for 5 min the pellets were air-dried and dissolved in water. The concentrations of RNA were measured using NanoDrop 2000 spectrophotometer. 1000 ng of each sample was used to synthesize complementary DNA. Total RNA was subjected to reverse transcription by Maxima H Minus Reverse Transcriptase (EP0753, Thermo Fisher Scientific) in the presence of 10 mM dNTP mix (R0191, Thermo Fisher Scientific) and oligo-(dT) (Biomers, Germany) was used to catalyze reverse transcription. qPCR was performed with Lightcycler 480 SYBR Green I Master mix (04887352001, Roche Diagnostics) using for measurements Lightcycler 480 Real-Time PCR System (Roche). The mRNA levels of the genes of interest were normalized to the mRNA levels of β-actin/GAPDH in corresponding samples.

### Primary cultures of sympathetic neurons and microinjection

Culture of mouse superior cervical ganglion (SCG) sympathetic neurons and microinjection of these neurons was performed as described earlier^81^. Briefly, the neurons of postnatal day 1–2 NMRI strain mice were grown 6 DIV on polyornithine-laminin (P3655 and CC095, Sigma-Aldrich)–coated dishes or glass coverslips with 30 ng/ml of 2.5 S mouse NGF (G5141, Promega) in the Neurobasal medium containing B27 supplement (17504044, Invitrogen). The nuclei were then microinjected with the expression plasmid for CDNF or CDNF mutants together with a reporter plasmid for enhanced green fluorescent protein (EGFP), at concentration of 10 ng/µl in each experiment. For protein microinjection, recombinant CDNF, or CDNF mutant proteins in PBS at 200 ng/µl were microinjected directly into the cytoplasm together with fluorescent reporter Dextran Texas Red (MW 70000 Da) (D1864, Invitrogen, Molecular Probes) that facilitates identification of successfully injected neurons. Next day, TM 2 µM (ab120296, Abcam) was added and living fluorescent (EGFP-expressing or Dextran Texas Red-containing) neurons were counted three days later and expressed as percentage of initial living fluorescent neurons counted 2–3 hours after microinjection.

### Human iPS-cell-derived dopamine neurons and treatment with CDNF mutants

iCell® DopaNeurons (R1088, FUJIFILM Cellular Dynamics) were seeded according to FUJIFILM Cellular Dynamics user protocol on 96-well plates coated with poly-L-ornithine (P3655, Sigma-Aldrich) and laminin (L2020, Sigma-Aldrich). Equal volumes of cell suspension were plated onto the center of the dish. The cells were grown for 5 days in iCell Neural Base Medium 1+ iCell Neural Supplement B (M1010, M1029, FUJIFILM Cellular Dynamics). Then the cells were treated with 6-OHDA, 50 µM, (H4381, Sigma Aldrich) and recombinant protein CDNF mutants (100 ng/ml or 1 µg/ml), without any trophic factor or with GDNF (100 ng/ml) (P-103-100, Icosagen), as negative and positive control, respectively. After three days, the cells were fixed and stained with anti-tyrosine hydroxylase (TH) antibody (MAB318, Millipore Bioscience Research Reagents). Images were acquired by ImageXpress Nano automated imaging system. Immunopositive neurons were counted by CellProfiler software, and the data was analyzed by CellProfiler analyst software. The results are expressed as percentage of living neurons, compared with untreated neurons in the control wells.

### Neurite outgrowth analysis in human iPS-cell-derived dopamine neurons

Human iCell® DopaNeurons were seeded in 96-well plates and subjected to treatment with either CDNF and CDNF mutants (10 ng/ml and 100 ng/ml), GDNF (100 ng/ml) as a positive control or left untreated as a negative control. After five days of incubation, cells were fixed and stained with antibodies against MAP2 (AB5622, Sigma-Aldrich) and tyrosine hydroxylase (TH), as well as DAPI (D9542, Sigma-Aldrich) for nuclear visualization. Automated imaging of the stained cells was performed using the ImageXpress Nano system. Neurite outgrowth parameters, including total outgrowth, the number of processes, and branching points, were quantified using CellProfiler software, based on MAP2 immunostaining^82^. The total neurite length (in µm) was determined by measuring the MAP2-positive regions, while the number of neurons was estimated by identifying double-positive signals for MAP2 and DAPI. Neurite length per cell was calculated by dividing the total neurite length by the average number of neuronal cells^83,84^.

### Quantification and statistical analysis

GraphPad Prism 9.0 software was used for statistical analysis. Statistical tests and sample sizes are indicated in the figure legends. p < 0.05 was considered as statistically significant.

