## Supplementary figures and images for "CDNF rescues human iPSCs-derived dopamine neurons through direct binding to unfolded protein response sensors PERK and IRE1α"

### Supplemental Figures S1-S6

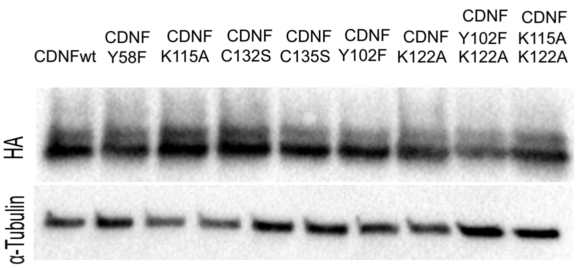

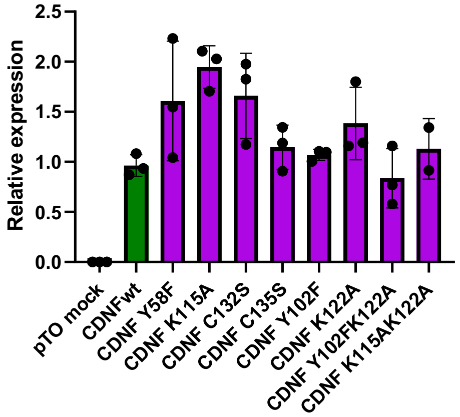

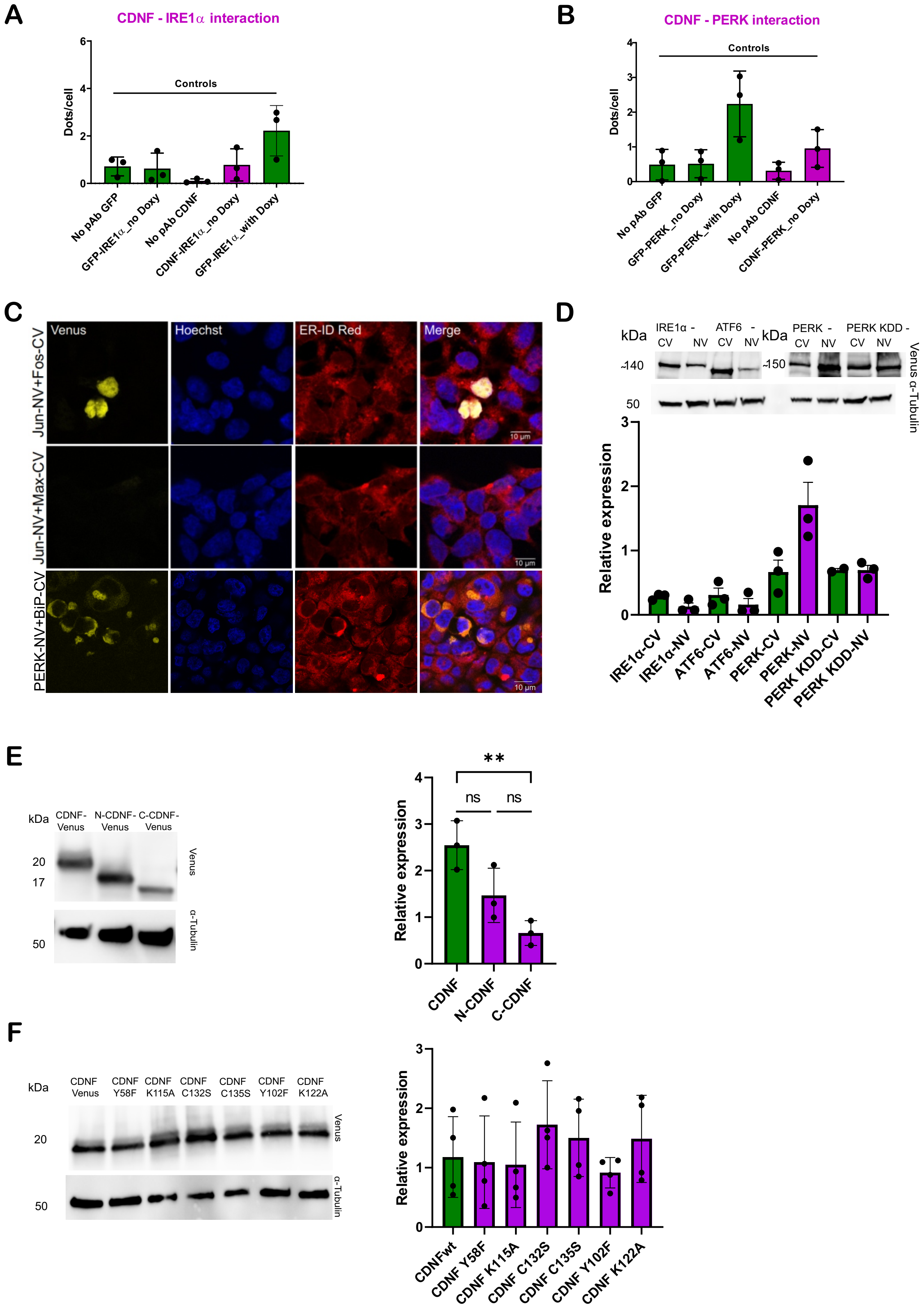

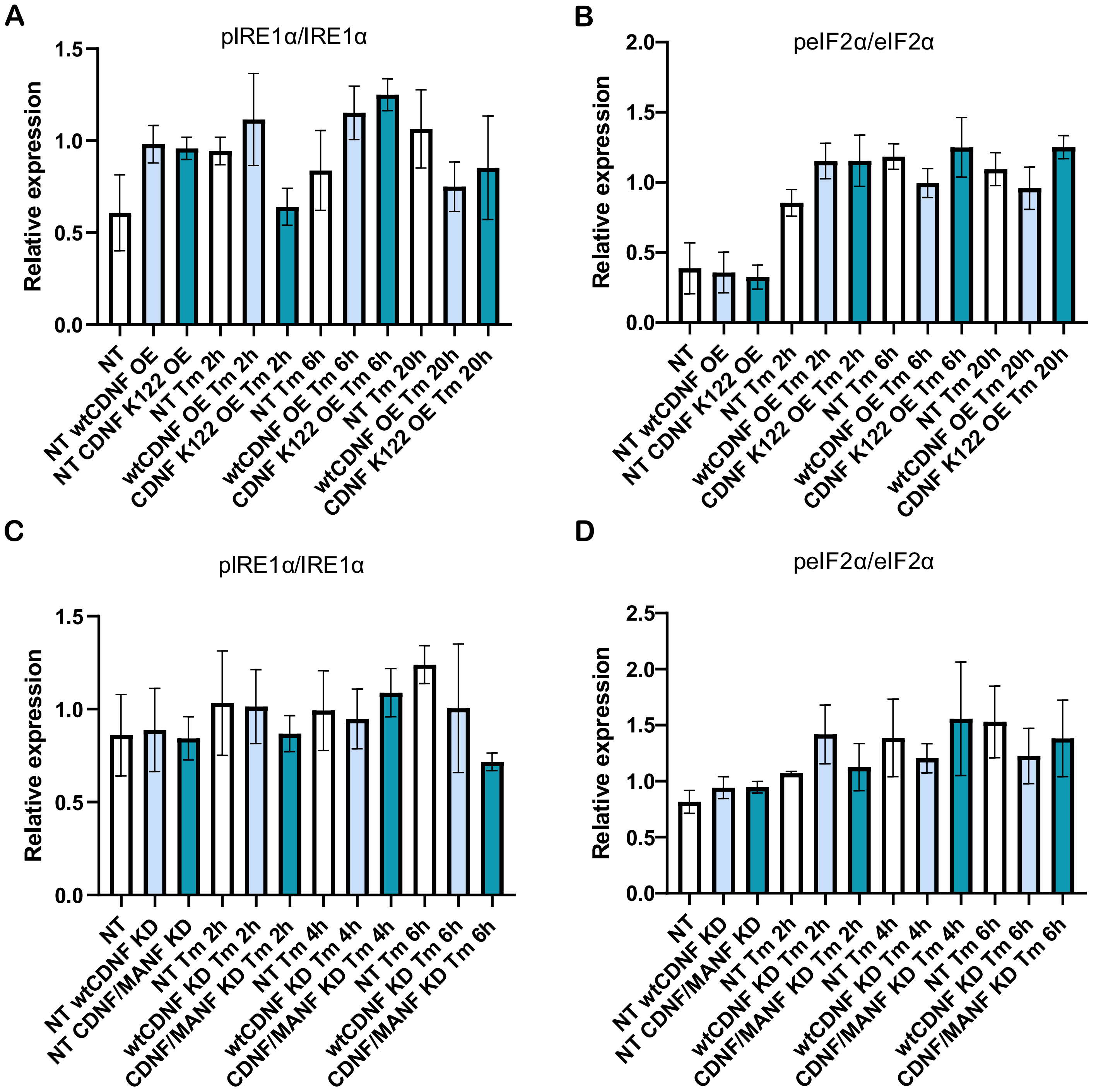

Figure S4

**A**

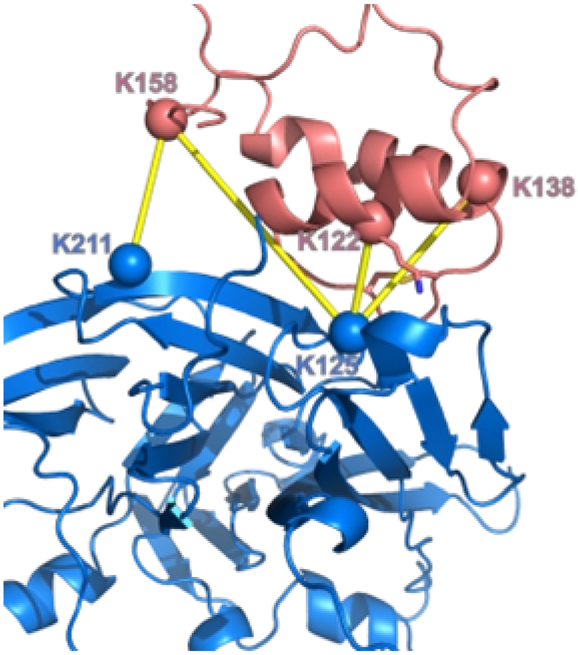


**B**

**CDNF + PERK LD**

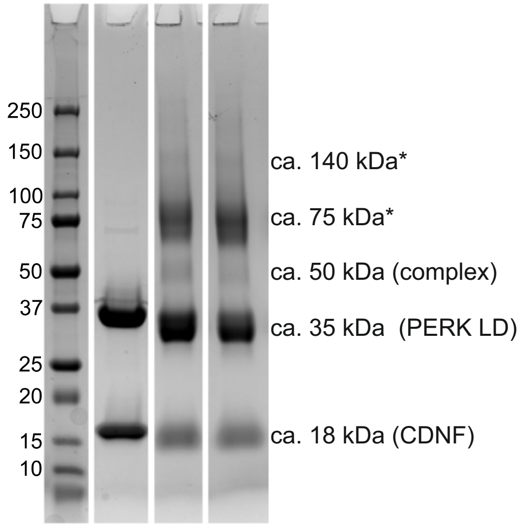

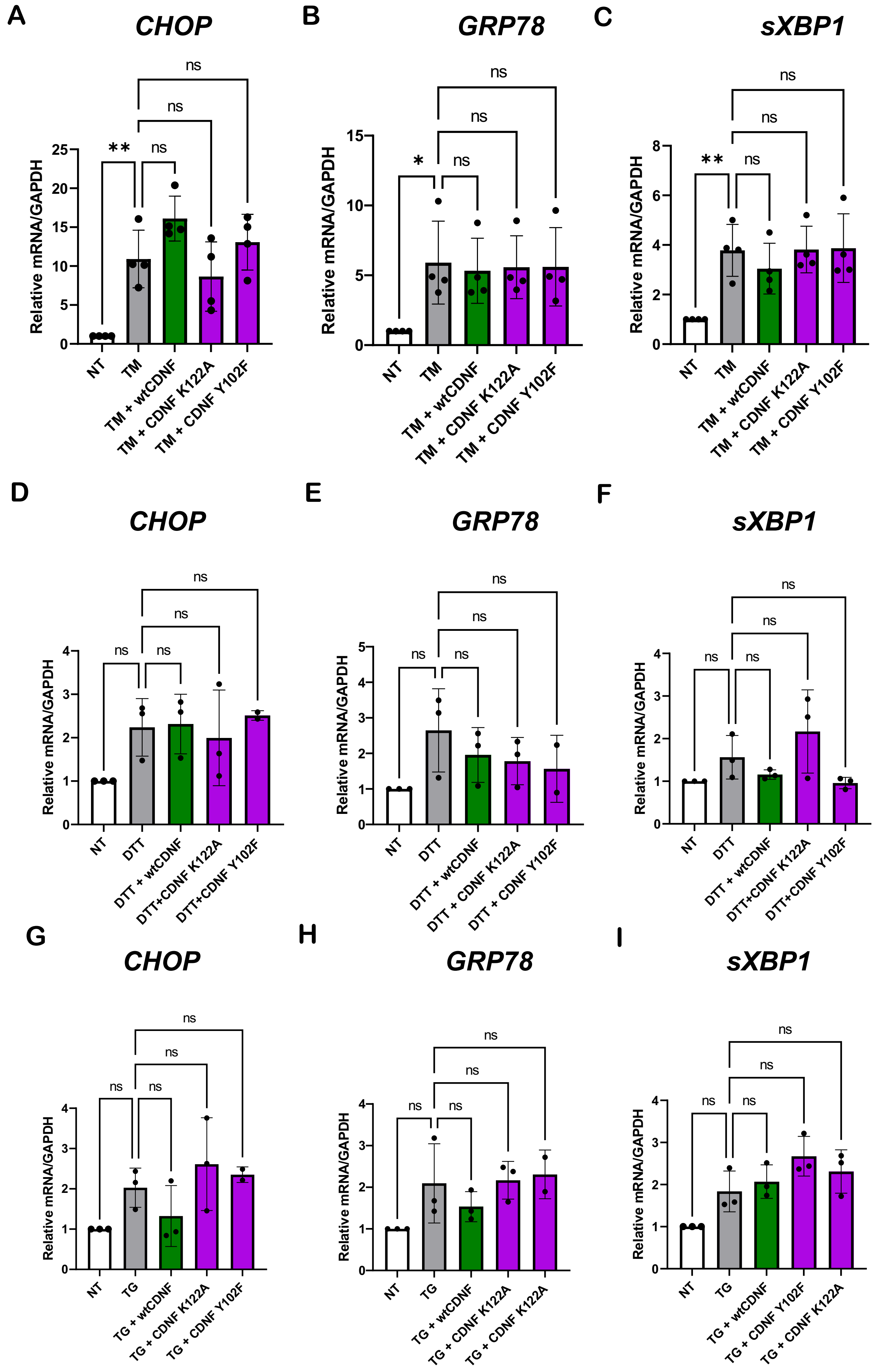

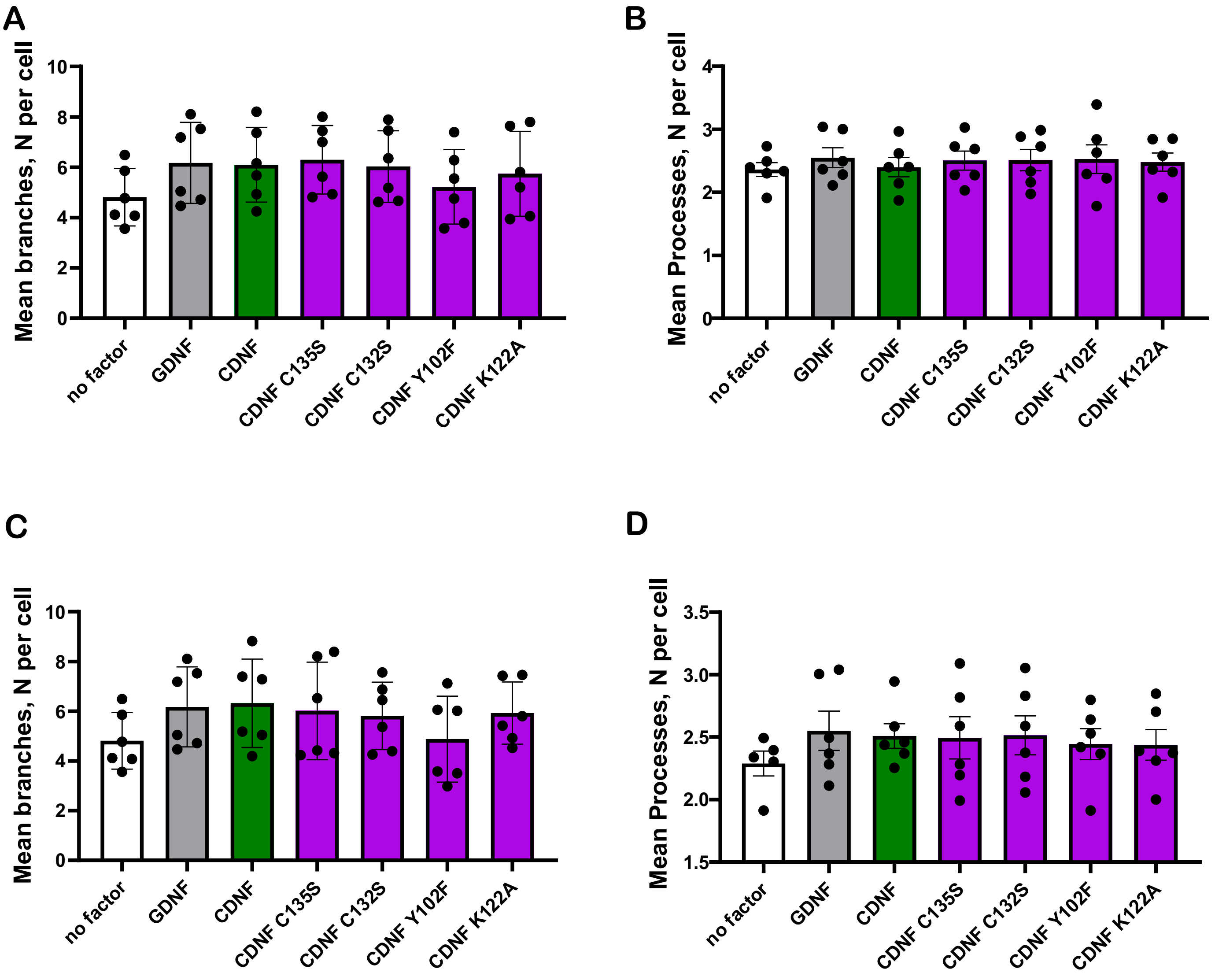
